# Data coherence over data volume drives generalisable genome-based prediction of microbial carbon utilisation

**DOI:** 10.64898/2026.08.06.743333

**Authors:** Dileep Kishore, Priya Ranjan, Christopher Neely, Mikaela Cashman, Zahmeeth Sakkaff, William J. Riehl, Marcin P. Joachimiak, Janaka N. Edirisinghe, José P. Faria, Pamela Weisenhorn, Myra B. Cohen, Dale A. Pelletier, Robert W. Cottingham, Christopher S. Henry, Mitchel J. Doktycz, Adam P. Arkin, Paramvir S. Dehal

## Abstract

Microbial carbon utilisation shapes ecological niches, community assembly, and nutrient cycling, yet remains diflcult to predict from genomes despite well-characterised pathways. Genome-based machine-learning models often generalise poorly across datasets, a limitation commonly attributed to small or taxonomically biased training sets. We integrated binary growth phenotypes for 819 strains across 240 carbon sources from four datasets, focusing on 15 shared sources. Cross-dataset evaluation reduced balanced accuracy from 0.86 (within datasets) to 0.62, and restricting evaluation to in-clade relatives recovered only 0.03 of this loss. Because phylogeny alone could not explain the transfer gap, we restricted training to samples with coherent genotype–phenotype relationships, defined by agreement between experimental labels and annotated pathways. Concordance filtering doubled the number of carbon sources with reproducible genomic signals across datasets (from 6 to 12 of 15), often recovering known pathway genes, while matched random subsets remained at the unfiltered level. The smaller concordant set also approached the cross-dataset performance of models augmented with up to 1800 BacDive genomes per phenotype, demonstrating greater eflciency per sample. Concordance-trained models remained bounded specialists, but separated growers from non-growers among genomes the pathway rules called negative, mostly generalists with broad-specificity transporters. Model confidence made these bounds actionable, since abstention improved accuracy where models transferred and retraining on newly labelled low-confidence genomes gave the largest gains for weaker phenotypes. Collectively, these findings recast generalisation as a problem of label–mechanism agreement and demonstrate how genome-based predictions can be deployed within explicit applicability domains and improved through targeted phenotyping.

## Introduction

Microbial nutrient utilisation phenotypes are key determinants of ecological niches and underpin community assembly, nutrient cycling, and biotechnological applications [1–4]. Cultivation-based methods underlie most phenotype data but cannot characterise unculturable taxa and remain resource-intensive even for culturable organisms [5, 6]. Phylogeny has commonly served as a proxy for phenotype prediction; however, it is insufficient for novel organisms with limited phylogenetic context or for traits that are not phylogenetically conserved [7–9].

Advances in genomics offer a path to overcome these limitations [10, 11]. Genome-based predictions provide an alternative, enabling characterisation of both cultured and uncultured taxa while potentially uncovering the genetic mechanisms underlying phenotypic variation [12, 13].

Early comparative-genomics and machine-learning systems, including PICA and Traitar, learned broad panels of microbial phenotypes from gene or protein-family patterns and linked predictive features to traits [14–18]. More recent multi-trait models such as MICROPHERRET and BacDiveAI extend phenotype coverage or draw on high-quality curated training data, while metaTraits integrates culture-derived records with outputs from multiple genome-based prediction systems at genome-catalogue scale [19–21], and knowledge graphs such as KG-Microbe harmonise strain phenotype, growth condition, and taxonomy data across sources [22]. Mechanistic approaches such as GapMind and microTrait instead infer metabolic or ecological capabilities from curated genes, pathways, and logical rules [23–25]. For carbon utilisation specifically, Li et al. showed that statistical gene-content models can outperform automatically reconstructed constraint-based metabolic models, while also demonstrating strong phylogenetic confounding and sample-size dependence [7]. Together, these approaches establish the feasibility of genome-based phenotype prediction, but leave unresolved how measurements pooled across independent experimental collections affect cross-dataset transfer and whether learned predictors recur as mechanistically relevant features across datasets.

Machine learning (ML) models can capture the nonlinear relationships and high dimensionality common in genomic data [26–29]. However, their performance and generalisability are constrained by data limitations and evaluation challenges [30, 31]: limited and taxonomically biased training data can cause overfitting and poor applicability to diverse genomes [12, 30, 32]. Phylogenetic correlations among genomic features further confound identification of mechanistically relevant predictors [33], since taxonomically biased models primarily capture phylogenetic signal [7]. Increasing sample size can improve out-of-clade prediction [7], and training on high-quality curated phenotypes can improve broader trait prediction [20], though such gains are typically established within a single curated source rather than across independent collections. Furthermore, if genotype–phenotype mappings are of mixed quality they may dilute the mechanistic signal available to the model [28, 34]. This suggests that label–mechanism agreement in training data, rather than only feature-space artefacts [35], is an important constraint alongside sample size.

We tested this hypothesis using microbial carbon utilisation as a model system. Carbon-utilisation phenotypes sit near the boundary of what genome-derived features can capture. The core biochemical transformations of many catabolic pathways are well characterised and conserved, yet their genomic mapping is often ambiguous. Transporters and broad-specificity enzymes are difficult to annotate, divergent or uncharacterised genes can perform similar chemistry, and regulatory mechanisms are inconsistently conserved across phylogeny. We integrated binary growth outcomes for 819 microbial strains across 240 carbon sources from four independent experimental collections and developed phenotype-specific classifiers for 15 shared phenotypes.

Models trained on concordant samples, those whose measured growth matches their annotated pathways, act as specialists. On held-out concordant samples they generalise better across datasets; they also more often recover established pathway genes as top features and rescue some samples misclassified by GapMind, a pathway-completeness predictor. This specialisation comes at the cost of coverage: the discordant samples set aside are precisely those whose genotype–phenotype links fall outside curated pathway definitions, so the resulting models apply to pathway-consistent genomes rather than to the full diversity of genotype–phenotype relationships. Genome-based phenotype prediction is therefore constrained not only by training-data size but by whether the labels agree with the mechanism being learned. Accuracy is consequently conditional on the genomes a model was built to describe, rather than a single value reported per phenotype.

## Materials and Methods

### Carbon Source Utilisation Datasets

We obtained four binary growth datasets on defined carbon sources: ATLeaf [36], Biolog [37], Marine [38], and Populus [39], harmonised into the phenotype tables deposited on Zenodo [40] (assay details in Supplementary Text S1 and Table S1). These contributed 819 input strain records, of which 795 had a genome assembly available for annotation. Genome quality (CheckM2) and pangenome completeness were assessed as covariates rather than applied as filters (Supplementary Text S2; Supplementary Figure S4). We aligned phenotype and genomic-feature identifiers, yielding 780 matched strains eligible for feature-based analyses. Sample sizes varied by phenotype because not every strain was measured for every phenotype. Carbon source names were standardised and analyses were restricted to the 15 phenotypes shared across all four datasets. A complementary data-volume comparison for 13 phenotypes additionally drew training samples from a fixed BacDive snapshot [41] (Supplementary Text S11).

### Genome Annotation

KofamScan v1.3.0 [42] assigned KEGG Orthologs (KOs) using KOFAM profile hidden Markov models from the KEGG database [43]. GapMind [23, 24] predicted carbon-utilisation pathway completeness, scored 0 (incomplete) to 2 (high confidence); it was run from the PaperBLAST suite [44] with USEARCH [45] and HMMER [46] (Supplementary Text S3). Rapid Annotations using Subsystems Technology (RAST) [47] provided functional predictions, parsed to extract subsystem ontology terms.

### Phylogenetic Tree Construction

We assigned taxonomy with GTDB-Tk [48] (GTDB release v220) and placed genomes into the GTDB [49] reference tree using classify wf. We extracted the corresponding subtree and computed distances between tips, the sum of branch lengths along the connecting path, with ETE 3 [50]. Genomes lacking species-level assignments were placed at genus or family level.

### Baseline Predictors

We evaluated GapMind as a mechanistic baseline using two binary thresholds on pathway-completeness calls, strict (only “complete” positive) and permissive (“complete” and “likely complete” positive), with all other calls negative. A phylogenetic *k*-nearest-neighbour classifier assigned phenotypes by majority vote among the *k* = 3 closest training relatives. Null baselines comprised an identity model (predicting the most common training class) and a Bernoulli model (sampling from observed training class probabilities).

### Data Pre-processing and Feature Selection

KOFAM, GapMind, and RAST annotations were converted into binary presence/absence matrices (GapMind pathway scores retained integer values). Two unsupervised pre-processing filters were applied to the combined feature matrix across datasets: variance filtering (scikit-learn [51] VarianceThreshold at 0.01) and correlation filtering (Spearman *ρ* > 0.95, graph-based, removing the most connected features).

### Machine Learning Classifiers

We trained gradient-boosted tree classifiers using CatBoost [52] (v1.2.8), which performed as well as or better than a range of alternative model families (Supplementary Table S2). The CatBoost hyperparameters were tuned on performance across all phenotypes (1000 iterations, learning rate 0.03, tree depth 4, used unless stated otherwise; Supplementary Text S3). Random holdout evaluations used five train/validation/test partitions generated independently for each phenotype and approximately stratified by its binary growth label, targeting a 70:15:15 allocation (Figure 3A). Cross-dataset and phylogenetic evaluations used a single 85:15 train:validation partition on the non-test data.

### Evaluation Strategies

#### Random Holdout and Cross-Dataset Testing

Performance was evaluated by balanced accuracy (BA) under four scenarios: random holdout within the aggregated dataset; leave-one-dataset-out cross-dataset; and in-clade and out-of-clade phylogenetic splits. Exact-duplicate feature vectors were rare and did not produce train–test overlap in leave-one-dataset-out evaluation (Supplementary Text S3). Phenotype–dataset combinations with fewer than 10 minority-class test samples were excluded to prevent BA collapse; cross-dataset missing features were imputed as zero.

#### Phylogenetic Partitioning

We applied the out-of-clade partitioning algorithm of Li et al. [7], which selects phylogenetically separated test clades (12–18% of samples). For cross-dataset evaluation, an in-clade test subset was instead defined by retaining test genomes whose average-linkage cluster of the phylogenetic distance matrix contained at least two training genomes (Figure 3C; distances by split type in Supplementary Figure S2).

### Concordant and Discordant Sample Analysis

Samples were stratified by agreement between GapMind permissive-threshold predictions and experimental outcomes: concordant (GapMind matches the experiment) or discordant (GapMind contradicts it); 71.9% of labelled samples were concordant across the 15 phenotypes and four datasets.

### Feature Importance Analysis

Stable predictive features were KOFAM features that recurred among the top ten by mean absolute Shapley additive explanation (SHAP) value across repeated models (Supplementary Text S7). Biologically redundant KOs were grouped post hoc into SHAP-supervised clusters before features were compared across datasets; clustering did not affect model fitting, predictions, or performance metrics (Figures 4C and 5B; Tables 1 and S3).

**Figure 1:**
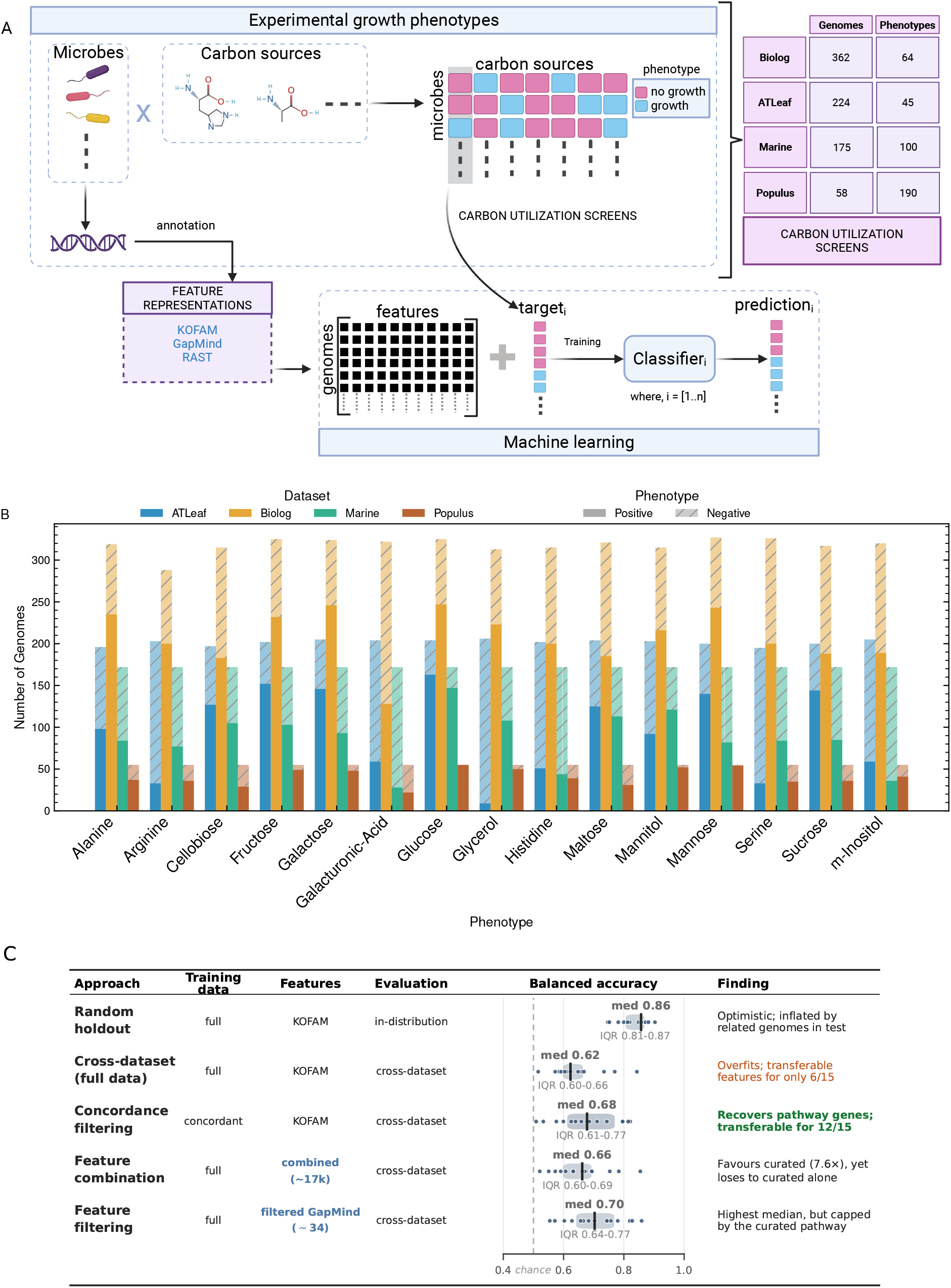
Workflow for genome-based carbon utilisation prediction, and the analysis variants evaluated in this study. (A) Overview of data collection, genome annotation, and machine learning workflow. We collected growth phenotypes (growth vs. no-growth) of microbial isolates across 240 carbon sources from four datasets (ATLeaf, Biolog, Marine, and Populus). Analyses focused on the 15 carbon sources shared across all four datasets. Genome sequences were annotated using RAST, KOFAM, and GapMind, and the resulting feature matrices were used in training the classifiers. The table in panel A gives the strain and carbon-source counts of each source dataset (Supplementary Table S1). (B) Phenotypic distribution for the 15 shared carbon sources across the four datasets. The bar plot displays sample counts and class distributions for each phenotype. Solid bars indicate growth (positive) samples, and patterned bars indicate no-growth (negative) samples. (C) Summary of the analysis variants evaluated in this study, showing for each its training data, feature set, evaluation, balanced accuracy, and a one-line finding. Bars span the interquartile range across phenotypes, the vertical tick marks the median, and dots show the individual per-phenotype values. The combined feature set concatenates the binarised RAST subsystem, KOFAM, and GapMind matrices (approximately 17,000 features). The 7.6× for feature combination is the rate at which curated GapMind terms appear among the top-ranked features of the combined-feature models relative to their share of that matrix; “loses to curated alone” refers to the phenotype-filtered set (last row). Panel A created in BioRender. Kishore, D. (2026) https://BioRender.com/tz7t8n7. **Alt text:** Panel A: workflow schematic linking microbial isolates and carbon sources to a growth or no-growth matrix, then to genome annotation with KOFAM, GapMind, and RAST and to classifier training, beside a table of strain and carbon-source counts for the ATLeaf, Biolog, Marine, and Populus datasets. Panel B: stacked bar chart of positive and negative sample counts for each phenotype in each of the four datasets. Panel C: table-style chart with one row per analysis variant, each row plotting balanced accuracy on a shared axis from 0.4 to 1.0 with a dashed line at 0.5, drawn as an interquartile-range bar with the median marked and one dot per phenotype. Medians and interquartile ranges are 0.86 and 0.81 to 0.87 for random holdout, 0.62 and 0.60 to 0.66 for cross-dataset full-data training, 0.68 and 0.61 to 0.77 for concordance filtering, 0.66 and 0.60 to 0.69 for feature combination, and 0.70 and 0.64 to 0.77 for feature filtering. A final column gives a one-line finding for each variant.

To establish baselines, we evaluated GapMind, a mechanistic tool predicting growth from pathway completeness [23, 24, 53], and a phylogeny-based nearest-neighbour classifier (*k* = 3; Methods). GapMind exposes substrate-specific pathway steps, so the same mechanistic framework provides a baseline and defines genotype–phenotype concordance. GapMind achieved modest BA across the 15 shared phenotypes (strict threshold: 0.66 ± 0.04; permissive: 0.70 ± 0.08; Figure 2A). The phylogeny-based approach performed well in random holdout (0.72–0.87) but poorly on phylogenetically distant samples (out-of-clade: 0.46–0.63; Figure 2B), approaching null-model levels and indicating that phylogenetic relatedness alone is insufficient across diverse lineages [8, 54–57].

**Figure 2:**
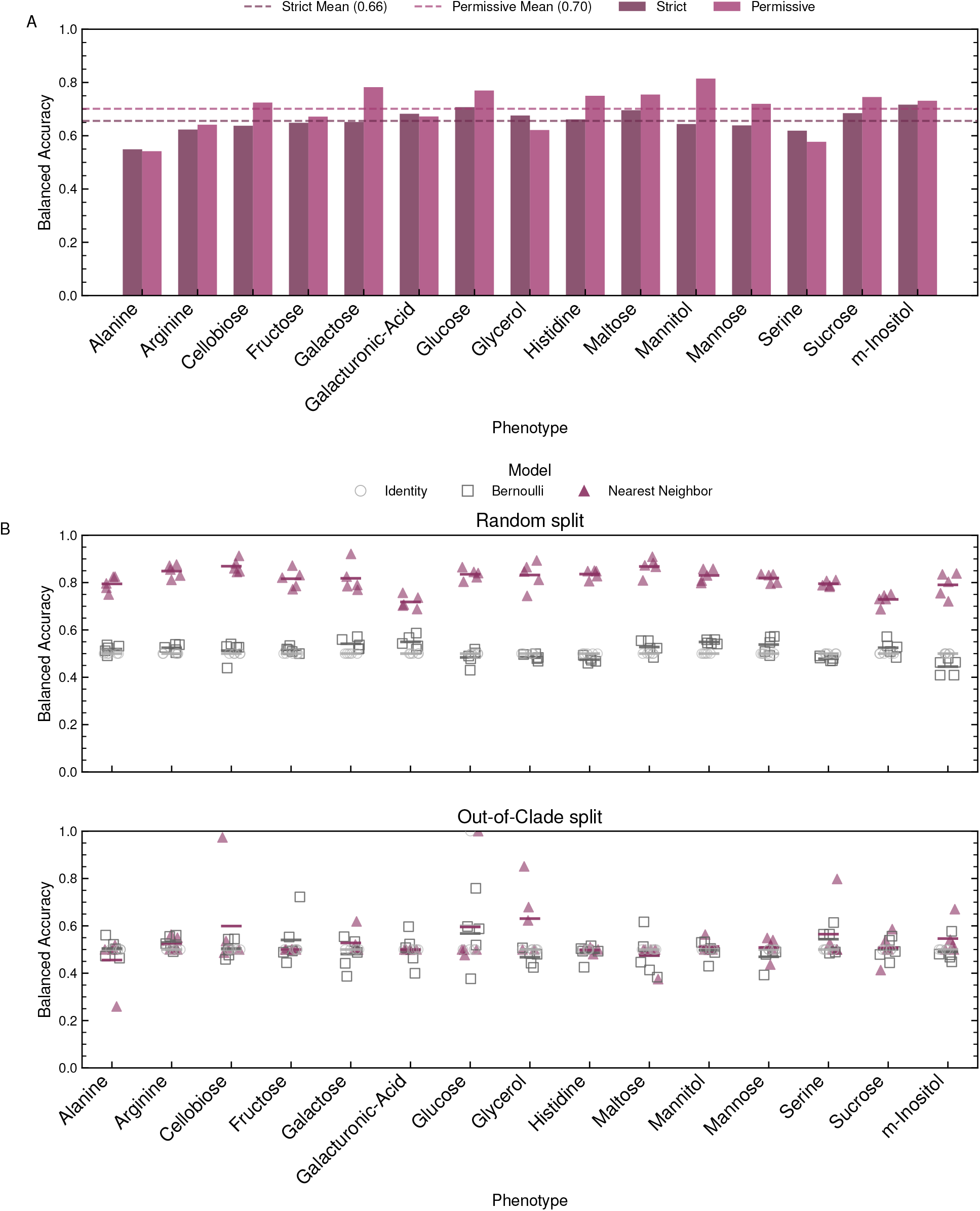
Neither curated pathway rules nor phylogenetic similarity predicts growth reliably. (A) Bar plot showing balanced accuracy for GapMind predictions across 15 phenotypes. GapMind predictions were compared with experimental outcomes using both strict (dark magenta) and permissive (magenta) pathway completeness thresholds. Dashed lines show the respective mean values. (B) Dot plot showing balanced accuracy for phylogeny-based nearest-neighbour (*k* = 3) predictions across 15 phenotypes under two evaluation scenarios: random split (top panel) and out-of-clade split (bottom panel). Circles and squares indicate null model performance (Identity and Bernoulli baselines) and triangles indicate nearest-neighbour performance. Each phenotype contributes five splits per model, and the short horizontal bar within each group marks the mean of those five values. **Alt text:** Panel A: grouped bar chart of GapMind balanced accuracy for 15 phenotypes under strict and permissive thresholds, with dashed mean lines near 0.66 and 0.70. Panel B: two dot plots of nearest-neighbour, Identity, and Bernoulli balanced accuracy under random and out-of-clade splits, with a short horizontal bar at each group mean. Values in the lower plot cluster near 0.5.

**Figure 3:**
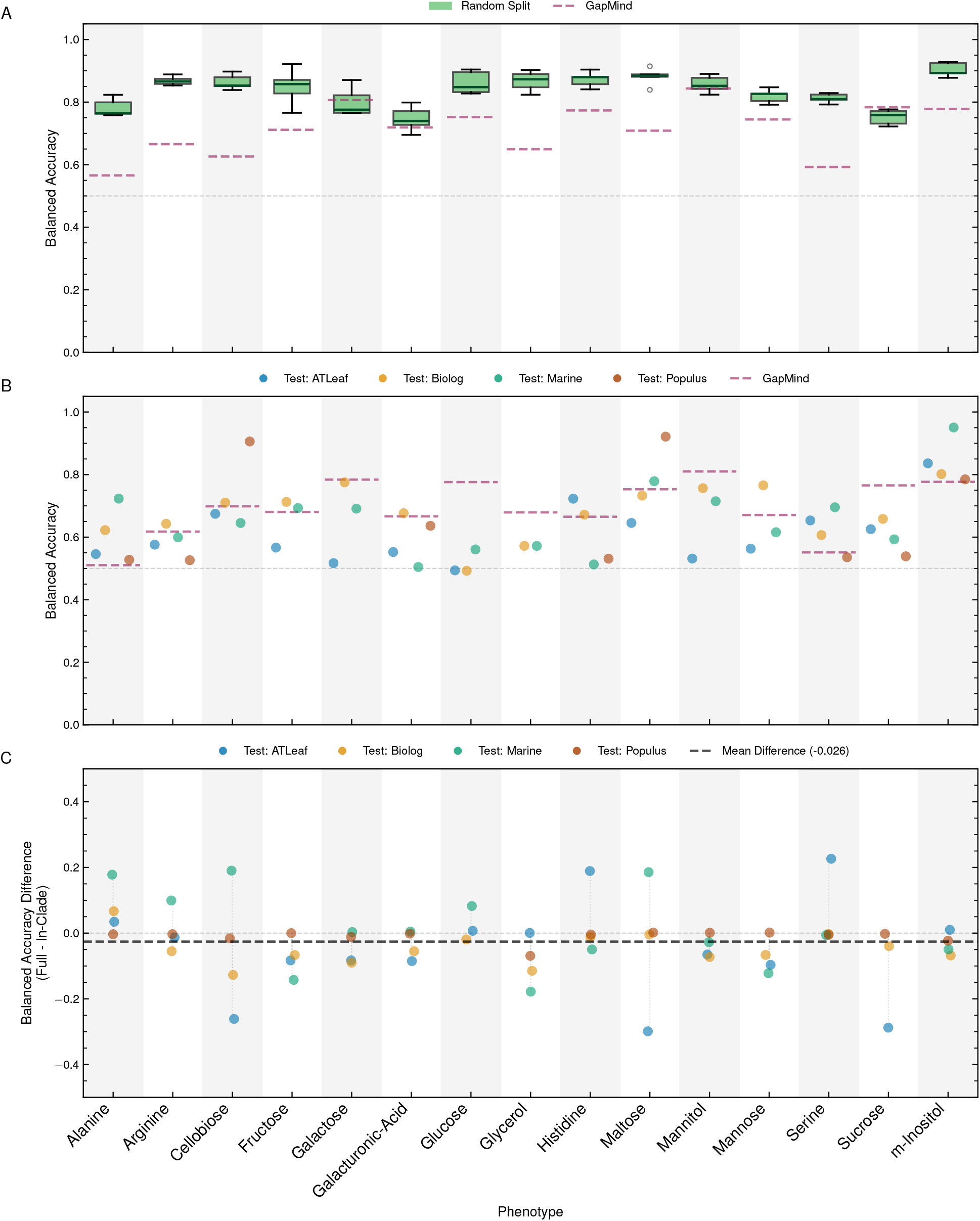
ML models are accurate on in-distribution genomes but lose that accuracy on unseen datasets, and phylogenetic distance does not explain the loss. (A) Box plot showing balanced accuracy under repeated random holdout evaluation for gradient-boosted tree classifiers (using KOFAM features) across 15 phenotypes. Each phenotype contributes five values, one per data-split. Boxes show the median and interquartile range. The dashed segment over each phenotype indicates that phenotype’s mean GapMind permissive-threshold balanced accuracy on the test set across splits. (B) Dot plot showing cross-dataset balanced accuracy when training on three datasets and testing on the fourth held-out dataset. Colours represent different test datasets. The dashed segment over each phenotype indicates that phenotype’s GapMind permissive-threshold balanced accuracy (same threshold as panel A). (C) Difference plot comparing model performance between regular cross-dataset splits and phylogenetically constrained in-clade cross-dataset splits. Colours denote the held-out test dataset as in panel B, and the black dashed line marks the mean difference across phenotypes and splits. Grey dashed horizontal lines mark chance performance (0.5) in panels A and B and zero difference in panel C. **Alt text:** Panel A: box plots of random-holdout balanced accuracy for each phenotype, clustering between 0.7 and 0.9 and sitting above the dashed GapMind baseline for all but two phenotypes. Panel B: dot plot of cross-dataset balanced accuracy coloured by held-out test dataset, scattered around the GapMind baseline. Panel C: dot plot of the difference between full and in-clade cross-dataset accuracy, centred near zero with a mean of minus 0.026.

**Figure 4:**
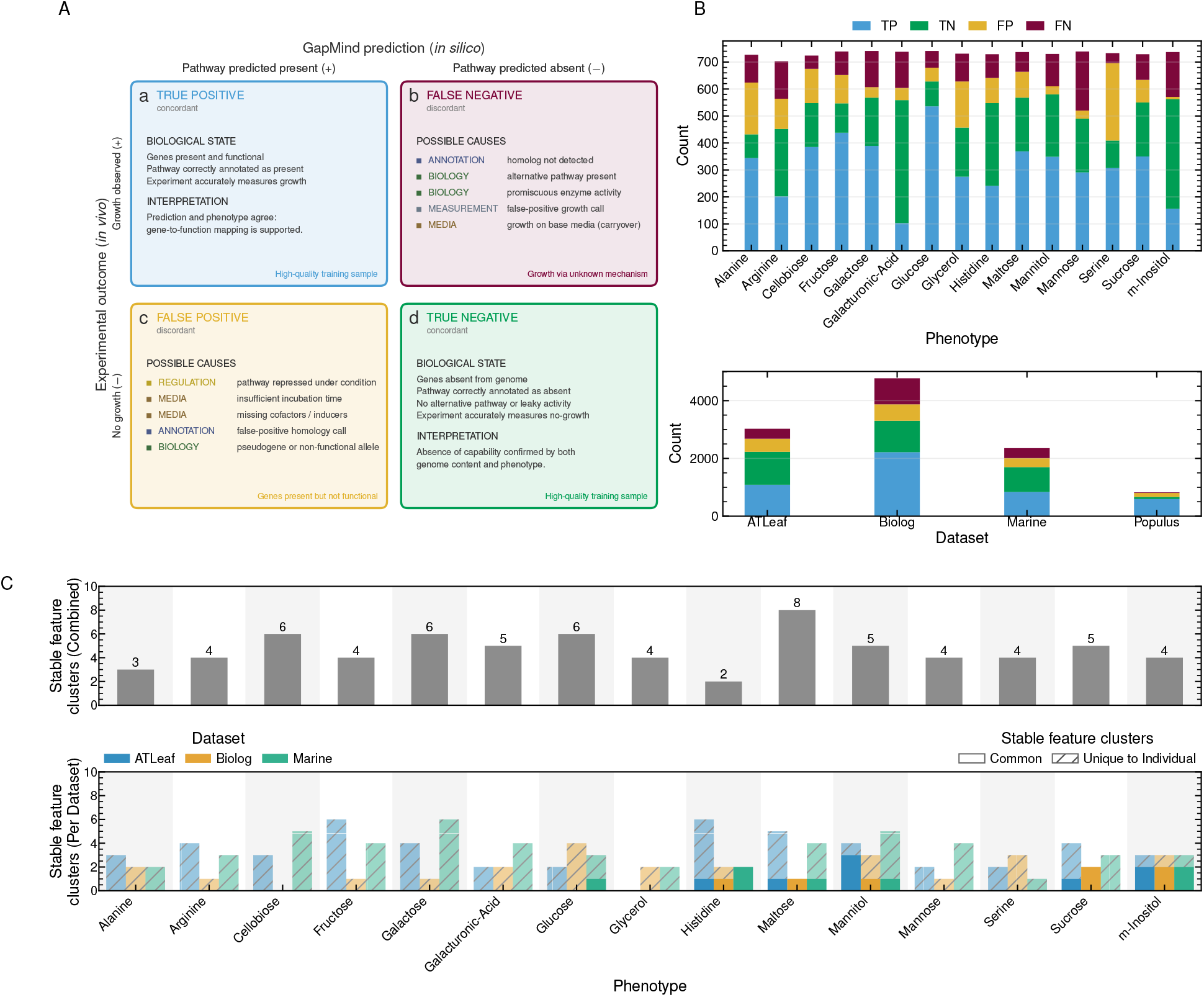
GapMind–experiment discordance is widespread, and ML models learn dataset-specific rather than shared features. (A) Conceptual framework contrasting concordant samples (where GapMind matches experiment) with discordant samples (where GapMind contradicts experiment). The concordant quadrants (a, true positive; d, true negative) give the underlying biological state and its interpretation, whereas the discordant quadrants (b, false negative; c, false positive) list possible causes colour-coded by category (annotation, biology, measurement, media, and regulation). (B) Stacked bar plots showing per-phenotype (top) and per-dataset (bottom) misclassification patterns. Each bar displays counts of true positives, true negatives, false positives, and false negatives. (C) Feature importance stability analysis using SHAP. Stable features are KOs appearing in ≥70% of model runs across 20 random seeds. These are grouped into stable feature clusters so that redundant KOs count as one (Methods). Top panel: number of stable feature clusters across the 20 random-seed runs when the model is trained on all four datasets combined. Bottom panel: cross-dataset cluster stability comparing the stable feature clusters from the model trained on the held-out dataset alone to those from the model trained on the three other datasets combined. Solid bars represent clusters shared between the held-out-alone and combined-of-three models and patterned bars represent clusters present only in the held-out-alone model. Populus is not shown as a held-out dataset because of its small sample size. **Alt text:** Panel A: two-by-two conceptual matrix mapping GapMind predictions against experimental outcomes, giving the biological state and its interpretation in the two concordant quadrants and colour-coded possible causes in the two discordant quadrants. Panel B: stacked bar charts of true-positive, true-negative, false-positive, and false-negative counts across fifteen phenotypes and four datasets. Panel C: bar charts of stable feature cluster counts per phenotype, for the model trained on all four datasets combined at the top, and for each held-out dataset at the bottom, where solid bars are clusters shared with the model trained on the other three datasets and patterned bars are clusters found only in the held-out-alone model.

**Figure 5:**
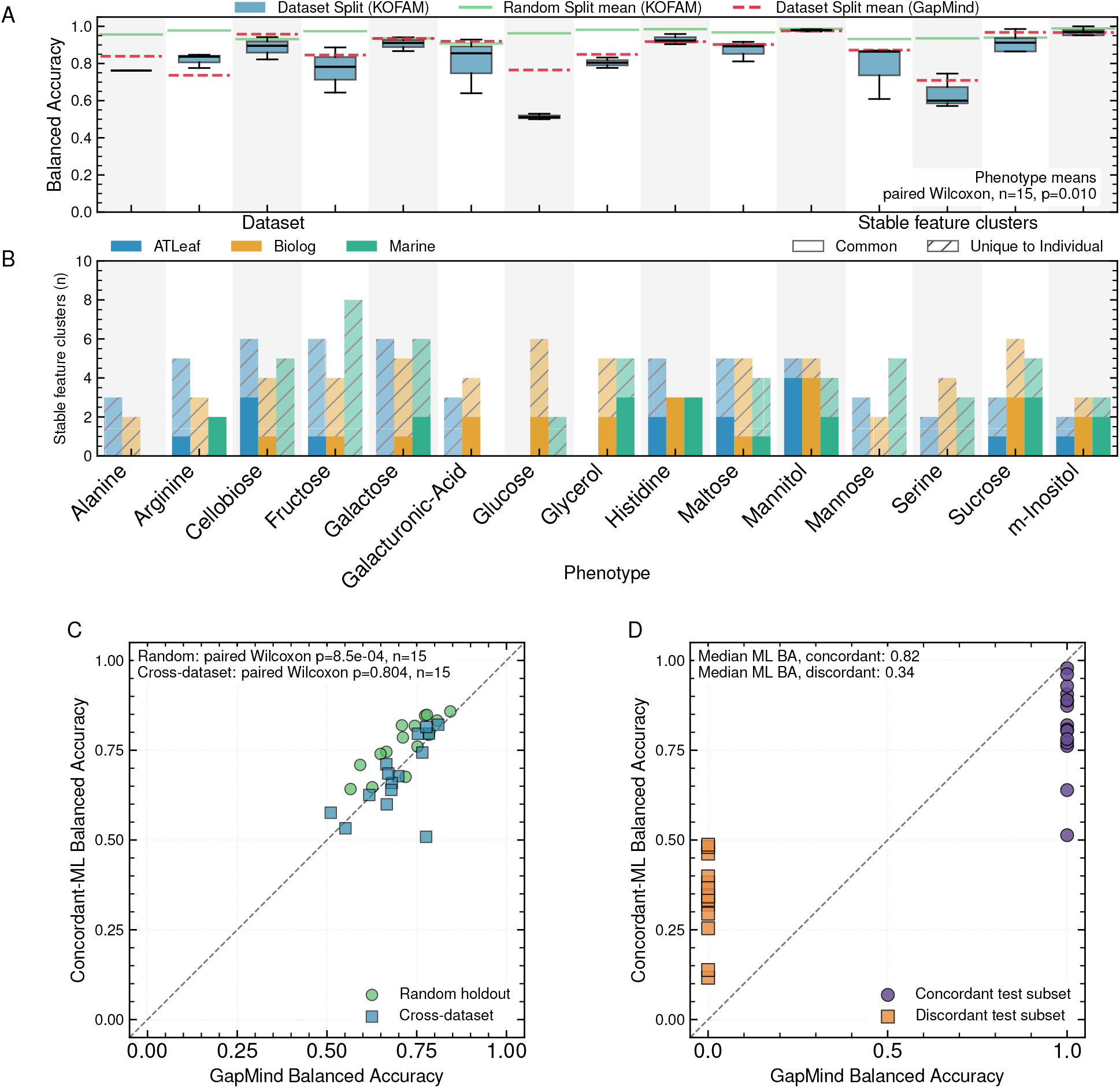
**Restricting training to concordant samples doubles the number of phenotypes with cross-dataset feature transfer and extends prediction beyond the subset GapMind classifies correctly**. All panels show models trained on GapMind-concordant samples with KOFAM features. (**A**) Balanced accuracy on held-out concordant samples, the easier subset with discordant cases excluded (panel C shows the full held-out test), under random holdout (green line) and cross-dataset evaluation (blue boxes). Red dashed lines mark a rule-reconstruction reference obtained by retraining on the GapMind step features whose completeness rule defines the concordant labels (Methods). Boxes show the median and interquartile range across the one to four leave-one-dataset-out splits retained per phenotype; outliers are not shown. Annotation: paired Wilcoxon *p*-value comparing the KOFAM and GapMind-feature phenotype means across phenotypes. (**B**) Cross-dataset stability of SHAP-supervised redundancy clusters between models trained on the three non-held-out datasets (Populus excluded) and on the held-out dataset alone. Each phenotype carries up to one bar per held-out dataset. Bars are absent where the concordant test set failed the minority-class exclusion (alanine and galacturonic acid in Marine, glycerol in ATLeaf), giving 42 rather than 45 comparisons. Solid bars: shared clusters; patterned bars: held-out-only clusters. (**C**) Concordance-trained ML versus GapMind balanced accuracy on the full sample held-out test set, one point per phenotype mean. Green circles: random holdout; blue squares: cross-dataset. Dashed diagonal marks parity; points above indicate ML outperforming GapMind. Annotation: paired Wilcoxon *p*-values across phenotypes, reported separately for the random holdout and cross-dataset series. (**D**) Cross-dataset test data decomposed into GapMind-concordant (purple circles, 15 phenotypes) and discordant (orange squares, 14 phenotypes; *myo*-inositol has no discordant set passing the minority-class exclusion). GapMind balanced accuracy is fixed at 1 on the concordant subset and 0 on the discordant subset by construction, so each series forms a vertical strip. Annotation: median ML balanced accuracy on each subset. **Alt text:** Panel A: box plots of cross-dataset balanced accuracy for each of fifteen phenotypes, with a line for random-holdout performance and a red dashed rule-reconstruction reference. Panel B: stacked bar chart per phenotype showing shared and held-out-only stable feature clusters for the ATLeaf, Biolog, and Marine held-out datasets. Panel C: scatter plot of concordance-trained model balanced accuracy against GapMind balanced accuracy, with random-holdout and cross-dataset markers lying near the dashed parity diagonal. Panel D: the same cross-dataset comparison split by subset, with concordant points forming a vertical strip at a GapMind balanced accuracy of 1 and discordant points forming a strip at 0.

### Data Quality Filtering Strategies

Two additional filtering strategies were evaluated beyond concordant sample filtering, both applied to training and validation samples only, never to the held-out test set. Problematic-sample removal excluded universal non-growers with any GapMind growth prediction, universal growers with incomplete GapMind pathways, and the twenty most frequently GapMind-misclassified genomes. Confidence-based weighting combined phylogenetic neighbour agreement (training labels only), GapMind pathway completeness, and experimental outcome. Four score configurations spanning *w*_gap_ ∈ {0, 0.3, 0.4, 0.5} were evaluated, including a mechanism-free setting (*w*_gap_ = 0) (full formula and sample weights in Supplementary Text S9). All conditions in Figure 6B were evaluated on the same full cross-dataset held-out test set.

**Figure 6:**
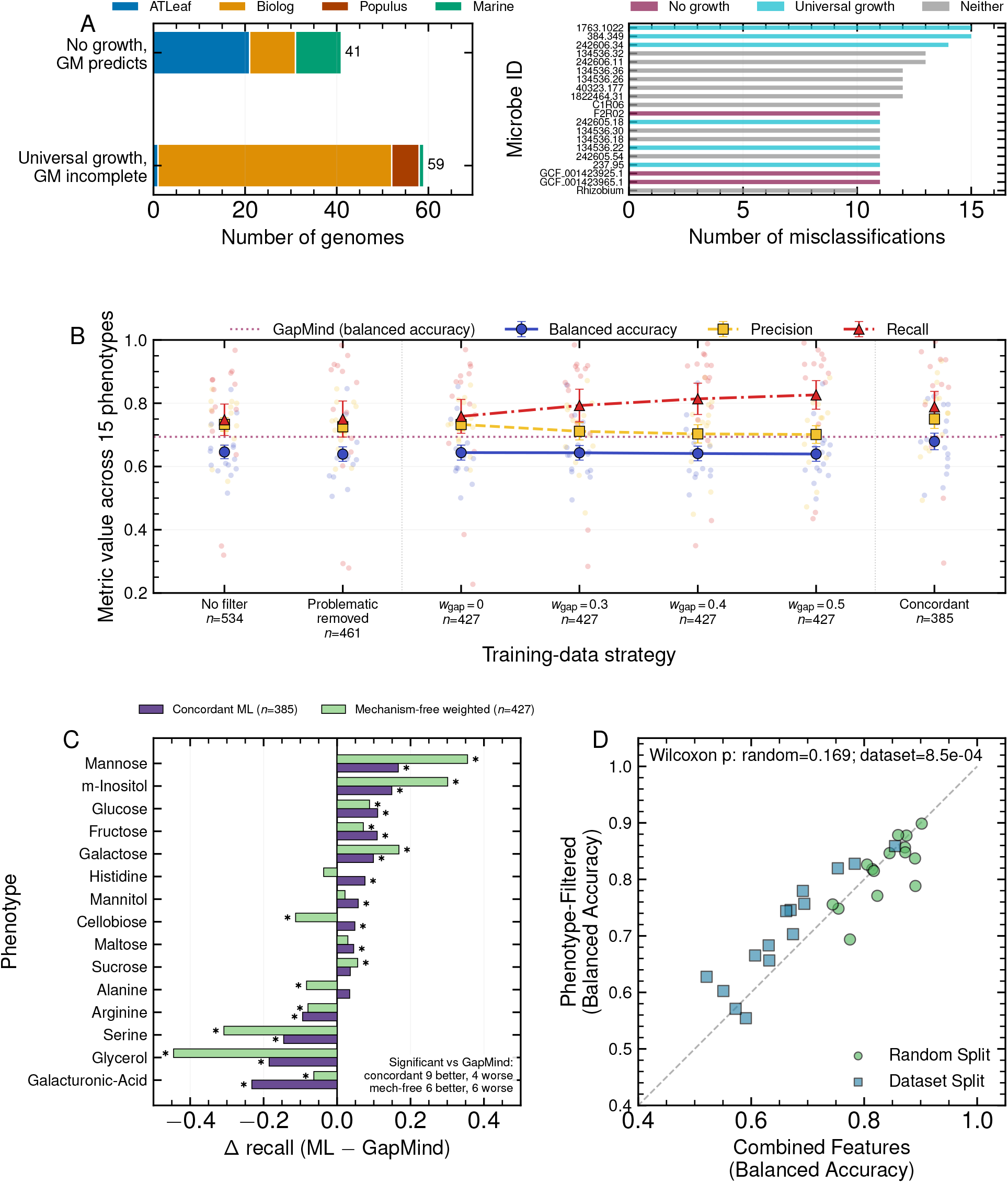
Confidence weighting shifts the precision-recall balance without changing accuracy, and more features do not beat fewer curated ones. (A) Left: counts of problematic genomes per dataset in two discordant categories: no observed growth despite at least one GapMind growth prediction, and universal observed growth despite incomplete GapMind pathways. Right: top-20 microbes ranked by number of phenotypes on which GapMind disagrees with the experimental label, coloured by membership in those categories from panel A. (B) Cross-dataset balanced accuracy, precision, and recall for soft-label weighting at *w*_gap_ ∈ {0, 0.3, 0.4, 0.5}, with three reference conditions plotted at the left and right: no-filter ML and problematic-sample removal (left), and concordance training (right). Large markers show the mean across the 15 phenotype-level values, error bars show the standard error of the mean across those values, and the small dots are the per-phenotype values. The mean train+val sample size per phenotype-split is annotated below each tick. The *w*_gap_ = 0 setting is the mechanism-free configuration, and all conditions are evaluated on the same full held-out cross-dataset test set. The dotted horizontal line marks the GapMind permissive-threshold rule-based baseline on the same test set. (C) Per-phenotype Δ recall (ML minus GapMind) on the same test set for concordance-filtered (purple) and mechanism-free-weighted ML (light green); phenotypes are sorted by concordant-ML Δ recall, and mean train+val sample sizes are given in the legend. Each bar is the difference in recall pooled over the genomes with observed growth in all four leave-one-dataset-out test sets, and asterisks mark the bars whose difference is significant at BH-adjusted *q* < 0.05 by an exact McNemar test on those same paired predictions. Annotation: the number of phenotypes each strategy significantly beats or loses to GapMind on. (D) Balanced-accuracy scatter under random holdout (green circles) and cross-dataset evaluation (blue squares) comparing combined (GapMind + KOFAM + RAST) and phenotype-filtered (the focal phenotype’s GapMind columns only) features. Points above the diagonal (equal performance, dashed) indicate that phenotype-filtered features outperform the combined set. The annotation reports split-specific paired Wilcoxon p-values across phenotype means. **Alt text:** Panel A: two horizontal bar charts, one counting problematic genomes in each dataset and one ranking the top twenty microbes by the number of phenotypes on which GapMind disagrees with the experimental label. Panel B: marker and error-bar plot of mean balanced accuracy, precision, and recall across the soft-label weighting sweep, with a dotted horizontal line marking the GapMind baseline and separate marker columns for the reference conditions at the left and right of the sweep. Panel C: bar chart of the per-phenotype change in recall relative to GapMind for the concordance-filtered and mechanism-free-weighted models, sorted by the concordance-filtered values, with asterisks marking the significant bars. Panel D: scatter plot of balanced accuracy for phenotype-filtered against combined features under random-holdout and cross-dataset evaluation, with a dashed diagonal marking equal performance.

### Sample Size Requirements

Training-data requirements were assessed for all 15 shared phenotypes by fitting full-data and concordance-trained KOFAM models to stratified subsets of 25, 50, 100, 200, and all available samples (three repeats) under random-holdout and cross-dataset evaluation, scored on full, concordant, and discordant test subsets. Subsets that retained only one class were skipped.

### Reliability and Experiment-Prioritisation Analyses

For full-data and concordance-trained models, prediction confidence was defined as max(*p*, 1 − *p*) and used as a ranking score; calibration was reported separately via reliability diagrams and expected calibration error. The receiver operating characteristic area under the curve (ROC AUC) for confidence-based discrimination of correct from incorrect predictions was pooled across phenotypes [51]. Risk–coverage curves retained genomes in decreasing order of confidence, scoring the retained subset by accuracy against the phenotype’s majority-class rate, since BA is undefined when the most-confident genomes share one true class. Mean confidence and the fraction of high-confidence predictions were Spearman-correlated with per-phenotype cross-dataset BA (Supplementary Text S8).

### Statistical Analysis

Paired comparisons of BA, precision, and recall across phenotypes used two-sided Wilcoxon signed-rank tests on phenotype-level means (scipy.stats.wilcoxon; Figures 3, 5, and 6). For the experiment-prioritisation comparison (Figure 7C,D), seeds were averaged within phenotype– held-out-dataset cells, with a paired test of the six phenotype means as a sensitivity analysis. Per-phenotype label-free aggregates were related to cross-dataset BA by Spearman rank correlation (scipy.stats.spearmanr; Supplementary Text S8). Thirteen primary tests (eleven Wilcoxon, two Spearman) were jointly adjusted by the Benjamini–Hochberg (BH) procedure at a false-discovery rate of 0.05; raw *p*-values are reported in the main text with adjusted *q*-values for borderline tests, and comparisons outside this family are unadjusted. Summary statistics are mean ± standard deviation across folds or repeats unless otherwise noted.

**Figure 7:**
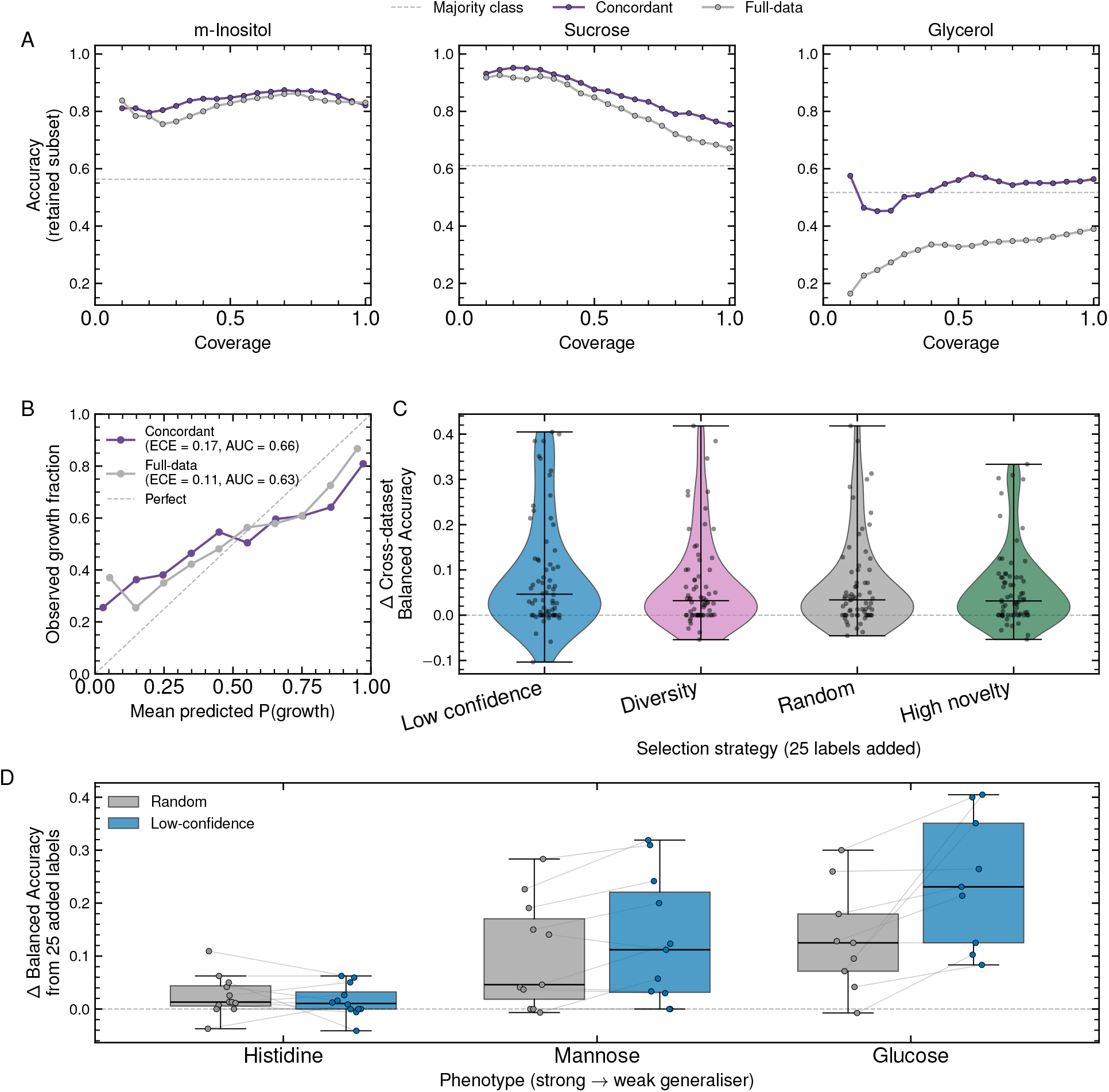
Model confidence supports selective abstention, and prioritising low-confidence genomes for follow-up is most effective where cross-dataset transfer is weakest. (A) Risk– coverage curves for three phenotypes spanning high, intermediate, and low cross-dataset balanced accuracy (m-Inositol 0.82, sucrose 0.74, glycerol 0.64), comparing the full-data model (grey) and the concordance-trained model (purple). Accuracy is shown on the retained subset as a function of coverage (the fraction of held-out genomes the model commits to) with the least-confident genomes abstained on first. The grey dashed line marks the phenotype’s majority-class rate, the accuracy reached by always predicting the more common outcome. Accuracy rather than balanced accuracy is plotted (Methods), so curves are read against the majority-class line rather than against 0.5. (B) Reliability diagram for both models on the cross-dataset held-out test set, pooled across phenotypes (10 equal-width probability bins), shows mean predicted probability of growth against the observed growth fraction within each confidence bin. The dashed diagonal marks perfect calibration, and the expected calibration error (ECE) and the pooled ROC AUC for confidence-based discrimination of correct from incorrect predictions are given in the legend. (C) Change in cross-dataset balanced accuracy after adding 25 held-out genome labels to the concordant training set, by label-free selection strategy (low confidence, diversity, random, high feature-novelty). Each strategy was run on six phenotypes with up to four held-out datasets and three seeds each, yielding 68 valid runs after excluding runs whose evaluation set contained a single class. Violins show the distribution across these runs (horizontal bar, median; vertical line, full range) and overlaid points show individual runs. (D) Per-phenotype gain from 25 randomly versus low-confidence-selected added labels for histidine, mannose, and glucose, ordered from strong to weak generaliser. These three come from the six phenotypes included in panel C and so differ from those in panel A. Boxes show the median and interquartile range, overlaid points show individual runs, and thin grey lines connect matched (held-out dataset, seed) runs across the two strategies (*n* = 12 for histidine, *n* = 11 for mannose, and *n* = 9 for glucose). **Alt text:** Panel A: three risk-coverage line plots, for m-Inositol, sucrose, and glycerol, each comparing the concordance-trained and full-data models against a dashed majority-class baseline. The m-Inositol curves are flat and well above the baseline, the sucrose curves rise steeply as coverage tightens, and for glycerol the concordance-trained curve hovers at the baseline while the full-data curve runs well below it and declines further at the tightest coverages. Panel B: reliability diagram in which both curves are flatter than the dashed diagonal, lying above it at low predicted probabilities and below it at high ones, indicating overconfidence. Panel C: violin plots with overlaid jittered points for the four selection strategies. Panel D: side-by-side box plots for histidine, mannose, and glucose comparing the random and low-confidence strategies, with thin grey lines connecting matched runs. The low-confidence boxes sit above the random boxes for mannose and glucose and level with them for histidine.

## Results

### Dataset overview and baseline predictions

To investigate carbon-utilisation prediction and generalisation, we harmonised four binary growth datasets (ATLeaf [36], Biolog [37], Marine [38], and Populus [39]) covering phylogenetically diverse strains on various carbon sources (819 input strain records, 795 with an available genome assembly, and 780 feature-linked strains; Supplementary Figure S1 and Table S1). We annotated genomes with RAST, KOFAM, and GapMind, trained classifiers on the resulting feature matrices (Figure 1A), and focused analyses on the 15 shared phenotypes (Figure 1B). Figure 1C provides a guide to the analyses that follow, with the BA of each variant shown alongside it.

### Machine learning models predict accurately within but not across datasets

To test whether genome-derived features could surpass these baselines, we trained gradient-boosted tree (CatBoost) classifiers on KOFAM annotations (∼5500 features; Methods), evaluating all models with BA.

Data were aggregated across all four datasets and assessed under four evaluation scenarios: repeated random holdout, cross-dataset (leave-one-dataset-out), and in-clade and out-of-clade phylogenetic splits (Methods).

In repeated random holdout evaluations (Figure 3A), the ML models achieved high BA (0.75–0.90) and outperformed the GapMind baseline (paired Wilcoxon across the 15 phenotypes, *p* < 0.001). Cross-dataset performance, however, decreased substantially (0.52–0.84; Figure 3B), indicating limited generalisation across experimental contexts [58]. The gap between regular and in-clade cross-dataset splits was minimal (∼0.03; Figure 3C), so phylogenetic distance alone does not explain the failure. Consistent with this, near-identical strains conflict on their growth label far more often across datasets than within a single dataset (Supplementary Text S12); this inconsistency limits cross-dataset transfer but does not prevent it, since most near-identical strains still agree. Out-of-clade counterparts showed similar trends (0.53–0.86; Supplementary Figure S3), consistent with prior observations that random holdout overestimates cross-context generalisation [7, 59].

### Cross-dataset failures reflect dataset-specific signals in aggregated training

To understand these failures, we first examined GapMind misclassification patterns across phenotypes and datasets (Figure 4A,B), which can arise from incomplete annotations, alternative routes, or regulatory mechanisms [23, 53, 60–62]. Phenotypes varied in GapMind accuracy and datasets varied in overall discordance, but neither showed a consistent directional bias, indicating that discordance does not arise from a single dataset-specific artefact (Figure 4B).

To probe generalisation failures, we performed SHAP-based feature-importance stability analyses, grouping KOs by SHAP-supervised redundancy (Methods) so biologically equivalent features count as a single signal (Figure 4C). Across random holdout evaluations of the aggregated dataset, at least one stable feature cluster was consistently identified, indicating recovery of recurring predictors when training data is sampled from the same pool (Figure 4C, top). By contrast, only six of fifteen phenotypes shared at least one common stable feature cluster between models trained on the three non-held-out datasets and the held-out dataset alone (Figure 4C, bottom; per-phenotype full-data counts in Table 1). Almost every held-out model still recovered stable clusters, but largely dataset-specific ones, outnumbering shared clusters roughly five to one (Figure 4C, patterned bars); the failure is therefore one of shared signal rather than learnable signal.

**Table 1:**
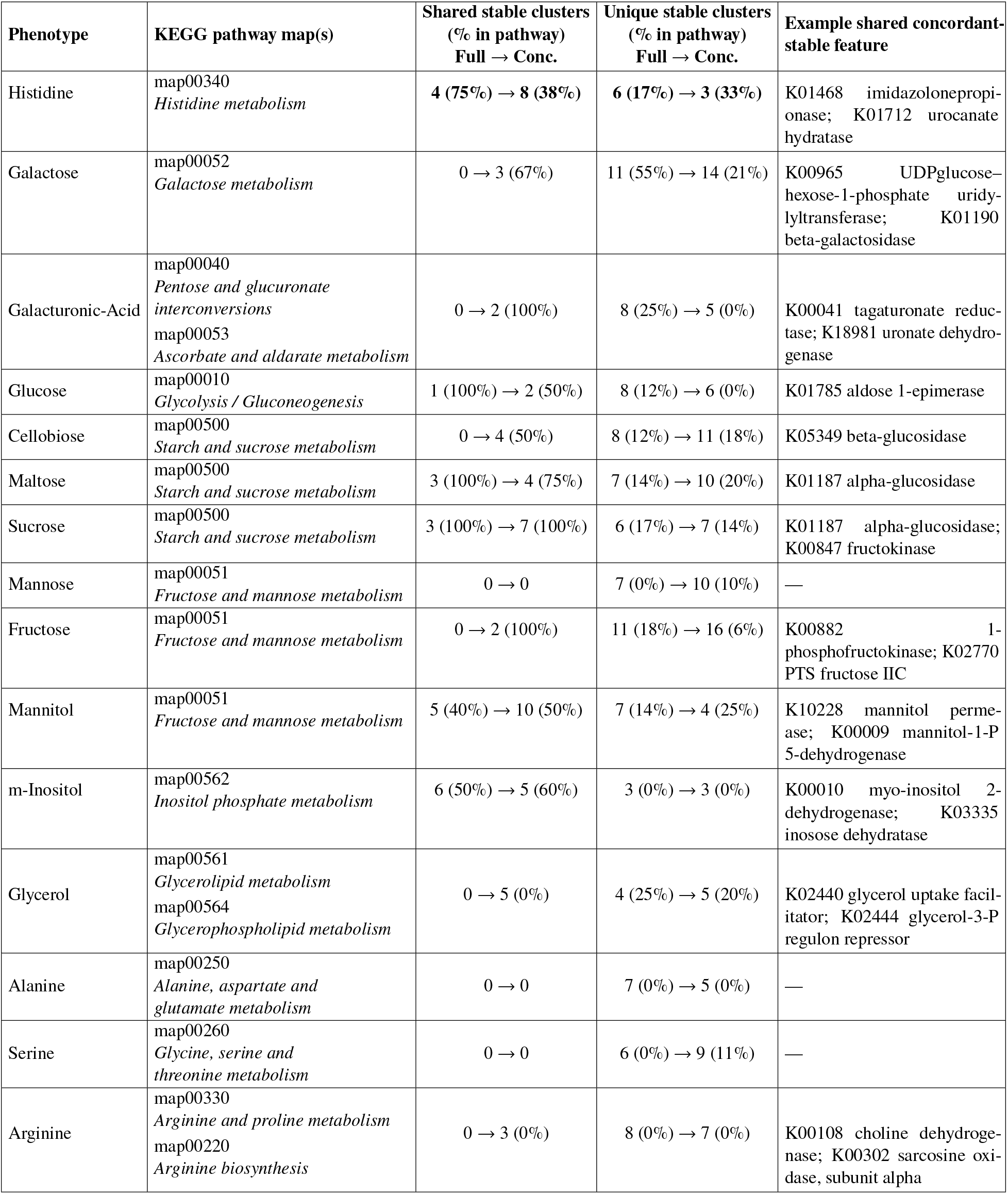
Concordance filtering recovers shared features across datasets for most phenotypes. For each phenotype, two models are compared: a three-dataset model trained on the three non-held-out source datasets pooled together, and a held-out-alone model trained on the held-out dataset alone. This pairwise comparison is repeated with each dataset in turn serving as the held-out one, and counts are summed across the retained held-out comparisons. The two columns labelled Full → Conc. report the same comparison under two training regimes applied to both models. Full uses all training genomes, while Conc. restricts training to GapMind-concordant samples. The shared stable-cluster count comprises SHAP-supervised redundancy clusters present in both the three-dataset and held-out-alone models (the same quantity plotted in Figure 5B), summed across the retained held-out comparisons, of which there are three per phenotype except where a comparison failed the minority-class exclusion (glycerol under full-data training; alanine, galacturonic acid and glycerol under concordant training). The unique stable-cluster count comprises those present only in the held-out-alone model on the same basis. The parenthetical (% in pathway) is the fraction of those clusters whose membership includes at least one KO on the row’s assigned KEGG reference pathway map. A cluster counts as pathway-resident when any of its KOs is on the map. The final example column reports one or two representative shared concordant-stable features for that phenotype, selected automatically and rendered with their KEGG descriptive names. An em dash (—) marks phenotypes with no shared cluster. This column is illustrative rather than exhaustive; the selection rules are given in Supplementary Text S4. Histidine is the worked example (numeric cells in bold); its per-comparison KO lists and the narrower unique-cluster module coverage (M00045 *histidine degradation*: 29% → 50%) appear in Supplementary Table S3 for full-data and concordant training. **Alt text:** Five-column table listing 15 phenotypes with their KEGG pathway map, shared stable cluster counts with parenthetical percent-in-pathway (Full to Conc.), unique stable cluster counts with parenthetical percent-in-pathway (Full to Conc.), and an example shared concordant-stable feature (enzyme, transporter, or regulator). Histidine row shows bold values rising from 4 (75 percent) to 8 (38 percent) shared clusters, with unique clusters falling from 6 (17 percent) to 3 (33 percent).

| Phenotype | KEGG pathway map(s) | Shared stable clusters<br>(% in pathway)<br>Full → Conc. | Unique stable clusters<br>(% in pathway)<br>Full → Conc. | Example shared concordant-stable feature |
| --- | --- | --- | --- | --- |
| Histidine | map00340<br>*Histidine metabolism* | 4 (75%) → 8 (38%) | 6 (17%) → 3 (33%) | K01468 imidazolonepropi-<br>onase; K01712 urocanate<br>hydratase |
| Galactose | map00052<br>*Galactose metabolism* | 0 → 3 (67%) | 11 (55%) → 14 (21%) | K00965 UDPglucose-<br>hexose-1-phosphate uridy-<br>lyltransferase; K01190<br>beta-galactosidase |
| Galacturonic-Acid | map00040<br>*Pentose and glucuronate*<br>*interconversions*<br>map00053<br>*Ascorbate and aldarate metabolism* | 0 → 2 (100%) | 8 (25%) → 5 (0%) | K00041 tagaturonate reduc-<br>tase; K18981 uronate dehydro-<br>genase |
| Glucose | map00010<br>*Glycolysis / Gluconeogenesis* | 1 (100%) → 2 (50%) | 8 (12%) → 6 (0%) | K01785 aldose 1-epimerase |
| Cellobiose | map00500<br>*Starch and sucrose metabolism* | 0 → 4 (50%) | 8 (12%) → 11 (18%) | K05349 beta-glucosidase |
| Maltose | map00500<br>*Starch and sucrose metabolism* | 3 (100%) → 4 (75%) | 7 (14%) → 10 (20%) | K01187 alpha-glucosidase |
| Sucrose | map00500<br>*Starch and sucrose metabolism* | 3 (100%) → 7 (100%) | 6 (17%) → 7 (14%) | K01187 alpha-glucosidase;<br>K00847 fructokinase |
| Mannose | map00051<br>*Fructose and mannose metabolism* | 0 → 0 | 7 (0%) → 10 (10%) | — |
| Fructose | map00051<br>*Fructose and mannose metabolism* | 0 → 2 (100%) | 11 (18%) → 16 (6%) | K00882 1-phosphofructokinase; K02770<br>PTS fructose IIC |
| Mannitol | map00051<br>*Fructose and mannose metabolism* | 5 (40%) → 10 (50%) | 7 (14%) → 4 (25%) | K10228 mannitol perme-<br>ase; K00009 mannitol-1-P<br>5-dehydrogenase |
| m-Inositol | map00562<br>*Inositol phosphate metabolism* | 6 (50%) → 5 (60%) | 3 (0%) → 3 (0%) | K00010 myo-inositol 2-dehydrogenase; K03335<br>inosose dehydratase |
| Glycerol | map00561<br>*Glycerolipid metabolism*<br>map00564<br>*Glycerophospholipid metabolism* | 0 → 5 (0%) | 4 (25%) → 5 (20%) | K02440 glycerol uptake facil-<br>itator; K02444 glycerol-3-P<br>regulon repressor |
| Alanine | map00250<br>*Alanine, aspartate and*<br>*glutamate metabolism* | 0 → 0 | 7 (0%) → 5 (0%) | — |
| Serine | map00260<br>*Glycine, serine and*<br>*threonine metabolism* | 0 → 0 | 6 (0%) → 9 (11%) | — |
| Arginine | map00330<br>*Arginine and proline metabolism*<br>map00220<br>*Arginine biosynthesis* | 0 → 3 (0%) | 8 (0%) → 7 (0%) | K00108 choline dehydroge-<br>nase; K00302 sarcosine oxi-<br>dase, subunit alpha |

Among the six phenotypes that did transfer, histidine is illustrative: KOFAM annotations consistently identified the main catabolic enzyme K01712 (urocanate hydratase) across datasets (Supplementary Text S4) [63]. Together, these analyses indicate that the aggregated training set pools many distinct genotype–phenotype mechanisms, mixing tractable samples with intractable ones (Figure 4C). We therefore hypothesised that restricting training to samples with more consistent genotype–phenotype relationships would let models recover transferable mechanistic features, using GapMind concordance to isolate this subset.

### Concordant training produces bounded specialist models

To test this hypothesis, we restricted training to concordant samples (GapMind predictions matching experimental outcomes) while keeping the feature space, architecture, and evaluation unchanged from Figure 3, so any gain reflects the training data alone (Supplementary Text S5; Supplementary Figure S5). Concordant samples likely overrepresent organisms using conserved pathways already captured in annotation databases, so the model is a specialist for mechanistically tractable cases rather than a recoverer of all valid utilisation strategies.

On held-out concordant test data (the easier subset, since discordant cases are excluded; full held-out test in Figure 5C), the concordance-trained model reached per-phenotype BA of 0.91–0.99 under random holdout (median 0.96) and median 0.82 under cross-dataset evaluation (Figure 5A). We then retrained the same models on the GapMind step features whose rules define concordance, giving a reference for how completely that rule is recovered from the step evidence itself (Figure 5A, red dashed lines). This reference exceeded the KOFAM models under cross-dataset evaluation (median 0.90 versus 0.82; paired Wilcoxon *p* = 0.01), with the gap below 0.1 for most phenotypes but largest for glucose (0.77 versus 0.51), whose substrate-specific transporter steps have no direct KOFAM equivalent (Supplementary Text S10).

Cross-dataset feature stability also improved under concordant training (Figure 5B; per-phenotype counts in Table 1). Twelve of fifteen phenotypes shared at least one stable cluster between three-dataset and held-out-alone models, up from six under full-data training (Figure 4C), and shared clusters per comparison rose 2.5-fold (1.31 versus 0.52 on the 42 matched comparisons; *p* < 0.001).

A size- and class-matched random-subset control remained at or below the full-data level (0.38 ± 0.03 versus 0.52), confirming the gain is specific to concordance, not subsetting (Supplementary Text S5). For histidine, concordant training yielded two urocanate-pathway enzymes shared in all three comparisons, added the histidine-utilisation repressor, and reduced dataset-specific features (Supplementary Text S4; Supplementary Table S3), consistent with the established co-regulation of *hut* catabolic genes [63]. For alanine, mannose, and serine, concordant cross-dataset accuracy was instead driven by broader carbohydrate-utilisation markers, a less specific but still in-domain mode of transfer (Table 1; Supplementary Text S7). Because concordant samples are those where GapMind’s pathway call already matched growth, this recovery is not independent of how the subset was chosen, although the stability of the same features across datasets does not follow from that selection.

To quantify what the concordance-trained KOFAM model adds over GapMind, we evaluated the full held-out test set (Figure 5C) and decomposed its cross-dataset portion by concordance status (Figure 5D). Under random holdout the ML model exceeded GapMind on 14 of 15 phenotypes (paired Wilcoxon *p* < 0.001; mean +0.05), but under cross-dataset evaluation the mean shift was undetectable across 15 phenotypes (paired Wilcoxon *p* = 0.80, BH-adjusted *q* = 0.87; Figure 5C). Under per-phenotype sample-level tests, sensitivity gains were matched by significantly lower specificity for 8 of 15 phenotypes, so the components largely cancel in BA. Stratifying the cross-dataset test set by concordance status separates two regimes (Figure 5D). Where GapMind already agrees with the measurement it is correct by definition, so the model’s median BA of 0.82 measures the feature-resolution gap discussed above rather than any gain over the rule. The informative cases are those where GapMind is wrong, and there the model’s binary predictions corrected few of them (BA 0.34), limited by how often it predicts growth at all. We therefore scored how well its predicted growth probability ordered genomes within each GapMind call, where the rule itself provides no ordering. It did so in both sets (pooled AUC 0.67), and concordant training helped specifically where GapMind had missed growth despite an incomplete annotated pathway (0.64 versus 0.53, BH-adjusted *q* = 0.003, 13 of 15 phenotypes), not where it had predicted growth that did not occur (*p* = 0.76). These recovered genomes were predominantly metabolic generalists carrying broad-specificity carbohydrate transporters and sugar-catabolism regulators, consistent with transport being the step GapMind most often failed to place (9 of 15 phenotypes; Supplementary Text S10).

These gains do not require large training sets: for most shared phenotypes, concordance-trained performance reached 90% of its all-sample value within 50 to 100 samples under both random-holdout and cross-dataset evaluation (Supplementary Text S6), although the phenotypes saturating earliest were also those with the lowest attainable accuracy. Adding up to ∼1800 BacDive genomes [41] to full-data training improved cross-dataset performance, but not significantly beyond a much smaller concordant set (13 phenotypes; Supplementary Text S11), and gave less reproducible feature selection (Supplementary Figure S7). Concordance filtering, however, requires a mechanistic predictor available only for well-characterised phenotypes and removes approximately 30% of samples, so we tested mechanism-free alternatives that use only phylogenetic agreement and experimental labels.

### Mechanism-free confidence weighting provides a recall-oriented alternative

Confidence weighting scored each tree-placed training and validation sample by whether its experimental label agreed with the labels of its nearest phylogenetic neighbours and with GapMind pathway completeness, down-weighting samples where these disagreed. The mechanism-free setting gave GapMind zero weight (*w*_gap_ = 0), and three comparison settings gave it 0.3 to 0.5 (Methods). A small number of problematic genomes, discordant with GapMind across many phenotypes, accounted for a disproportionate share of misclassifications (Figure 6A) and were tested as an additional filter.

Across the *w*_gap_ sweep, mean cross-dataset BA varied by less than 0.005 while recall rose and precision fell (Figure 6B), a shift towards positive calls rather than greater discrimination. Problematic-genome removal alone gave no measurable improvement. Concordant training had the highest mean BA (0.680 versus 0.644 for mechanism-free weighting), although the difference was not significant (paired Wilcoxon *p* = 0.23). Recall against GapMind is the informative axis here, because its errors are predominantly missed growth. Recovering those genomes indicates catabolic capability that the curated pathway does not represent. Mechanism-free weighting raised recall slightly over unfiltered ML with no change in BA and lower specificity. Concordant training produced significant phenotype-level recall gains over GapMind for more phenotypes (9 of 15 versus 6), although neither mean recall shift reached significance (Figure 6C). Recall gains were therefore consistent in direction but small on average, with the demonstrable advantage confined to the genomes GapMind calls incorrectly, where the model orders growth outcomes that the rule cannot (pooled AUC 0.67; Supplementary Text S10).

Curation can also be applied to the features rather than to the samples. Phenotype-filtered GapMind features (without correlation filtering; Supplementary Text S5) matched combined GapMind, KOFAM, and RAST features under random holdout but improved cross-dataset BA (paired Wilcoxon *p* < 0.001, BH-adjusted *q* = 0.003), and the combined-feature models still concentrated on curated GapMind terms, consistent with prior reports favouring biologically relevant over comprehensive feature sets [7] (Figure 6D). Sample filtering, sample weighting, and feature choice all act at training time and shift average performance; none indicates which individual predictions to trust. We therefore asked whether a model can flag which of its own predictions to trust from information available at inference time, without the experimental phenotype.

### Prediction confidence defines an applicability domain and prioritises follow-up experiments

Pooled across phenotypes, the model’s predicted probability was higher for predictions that proved correct than for those that did not, though only modestly (ROC AUC 0.66 concordance-trained, 0.63 full-data), and both models were overconfident under cross-dataset shift (Figure 7B). Consistent with prior reports [64], confidence therefore supports ranking and abstention rather than calibrated probability estimates. Abstaining on the least-confident predictions accordingly helped only where the model had transferable signal to concentrate (Figure 7A): for sucrose it raised retained-subset accuracy from 0.75 at full coverage to 0.88 at half coverage against a majority-class rate of 0.61, whereas for m-Inositol, already accurate across the full test set, it added little, and for glycerol it added nothing. Abstention can therefore define a practical applicability domain [65–67] but cannot substitute for a model that has not learned the phenotype.

Confidence did not, however, indicate which phenotypes a model would handle reliably: neither mean prediction confidence nor the fraction of high-confidence predictions correlated with cross-dataset BA (Supplementary Text S8). Since reliability cannot be certified per phenotype in advance, we asked instead which additional measurements would most improve a model, simulating the addition of 25 held-out genome labels to the concordant training set and retraining (Figure 7C). Selecting the least-confident genomes produced the largest mean gain in cross-dataset BA (0.09 versus 0.07 for random; *p* = 0.024, BH-adjusted *q* = 0.052), although the advantage was not consistent across the six phenotype means (*p* = 0.16), and selecting the most novel or diverse genomes did not beat random [68]. The advantage tracked how poorly the model already transferred (Figure 7D): largest for glucose, where low-confidence selection roughly doubled the gain from random labelling (0.24 versus 0.13), smaller for mannose, and absent for histidine, whose model already transferred well. Confidence therefore ranks usefully in two ways, flagging which predictions to withhold and which genomes to measure next, even though it does not reveal which phenotypes will transfer.

## Discussion

ML models that predict microbial phenotypes from genomic features generalise poorly across datasets [30, 58], a limitation typically attributed to limited sample size, taxonomic bias, or feature-space artefacts [7, 69, 70]. Across four binary growth datasets and the 15 carbon sources assayed in all of them, we identify label–mechanism agreement in the training set as a determinant of cross-dataset generalisation that complements the established roles of sample size, phylogenetic structure, and feature-space choice.

### Label–Mechanism Agreement Governs Cross-Dataset Transfer Within a Bounded Domain

Poor cross-dataset performance was better explained by mixed-quality training samples and dataset-specific correlations that did not transfer than by clade structure, which accounted for little of the failure (Figure 3C). Restricting training to samples whose experimental labels agreed with mechanistic predictions (concordance filtering) improved cross-dataset feature stability. Although the classifiers have no mechanistic knowledge, concordant training led them to converge across datasets on curated pathway genes (Figure 5B; Table 1).

Discordance has several coexisting causes [71, 72]: false negatives may reflect annotation gaps or unknown alternative pathways, whereas false positives may reflect regulatory repression or experimental artefacts such as differences in growth-assay modality across the source datasets, which static genomic features cannot capture (Figure 4A). The model correctly predicted growth for some of the discordant genomes, most of which were metabolic generalists carrying broad-specificity transporters and sugar-catabolism regulators. GapMind excludes such genes by design, since a broad transporter’s presence does not establish that it serves the focal substrate. Across the whole cross-dataset test set the concordance-trained model did not exceed GapMind, and on concordant genomes it cannot (by definition). The gain is therefore limited to the genomes GapMind gets wrong. Among those, the rule returns a single verdict and cannot tell them apart, whereas the model assigns higher growth probabilities to the genomes that do grow, most clearly where the annotated pathway looked incomplete (Supplementary Text S10). The model adds to the rule rather than replacing it, and the discordant genomes removed by filtering are the ones that carry this information.

Li et al. reported that datasets exceeding approximately 1000 genomes can enable out-of-clade prediction for some traits [7], whereas Yu et al. found that increasing sample size did not rescue generalisation under phylogenetically biased sampling [69]. Within our sample-size regime (up to 780 strains), the heterogeneity of training samples rather than their number alone drove shortcut learning, extending that second result from taxonomic to mechanistic heterogeneity. Curated data quality is known to improve trait prediction [20], but concordance is a criterion of a different kind. Discordant records are not necessarily erroneous, since a genome may grow through an uncharacterised route or fail to grow despite a complete pathway, so concordance removes mechanistic heterogeneity rather than error. Adding up to ∼1800 BacDive genomes per phenotype [41] improved cross-dataset performance over full-data training but not significantly beyond a roughly four-fold smaller concordant training set, and reduced feature reproducibility (Supplementary Text S11).

### Practical Deployment and Confidence-Guided Experimentation

Concordance filtering cannot be applied without a mechanistic predictor for the phenotype in question, which exists only for well-characterised pathways. One alternative is to weight samples by phylogenetic label agreement, which requires no pathway annotations and increased true-positive recovery, though with lower specificity and little BA change from unfiltered ML (Figure 6B,C) [73, 74]. When GapMind is available for every genome, a simpler option is to restrict a full-data model to the predictions that agree with it, which reached BA 0.77 on the 72% it retained (Supplementary Text S10). Explainable predictors whose rules are learned from harmonised phenotype data, rather than curated, are another route to such a check [75]. A second constraint is how much labelled data these models need, measured on cross-dataset tests since random holdout overestimates generalisation [59]. A small concordant training set is enough to indicate whether a phenotype is learnable from genomic features, since 50 to 100 concordant samples reached at least 90% of the all-available-sample performance for most shared phenotypes (Supplementary Figure S6). Because the earliest-saturating phenotypes were also the least predictable, further measurement is worth committing mainly where that early signal exists.

Committing that measurement effort and deciding whether to trust a prediction on a genome whose concordance is unknown require a signal that does not depend on GapMind, and model confidence provides one, flagging which predictions to withhold [65–67]. Among the strategies we compared for choosing genomes to label, selecting the least-confident ones produced the largest mean transfer gain, generally concentrated in the weakest phenotypes, whereas diversity and novelty-based selection did not beat random (Figure 7C,D). High-throughput phenotyping methods [38, 76–78] could scale this loop, directing new measurements towards low-confidence genomes and under-sampled clades in the weakest-transferring phenotypes rather than adding depth where performance has already saturated. At the phenotype level, however, no label-free metric we tested correlated with cross-dataset accuracy. Phenotype difficulty appears instead to be a property of the feature space rather than of the model, since confidence reflects agreement among the patterns a model has already seen and cannot register a mechanism that the annotations do not encode or that occurs too rarely to be learned.

### Limitations and Future Directions

Concordance-based training requires mechanistic tools such as GapMind, available only for well-characterised pathways, and mechanism-free weighting avoids that requirement only by trading specificity for recall while still needing experimental labels. Genome-scale metabolic models [79, 80] could extend concordance filtering to a much larger set of carbon substrates, though this would require curated or optimised reconstructions, since automatically constructed constraint-based models have been reported to predict carbon utilisation little better than chance for non-model organisms [7]. Filtering also extends the tradeoff between specialisation and coverage to taxonomy, favouring pathways in the annotation space and organisms with broadly conserved enzymes while excluding those with alternate or unknown enzymes, divergent pathway variants, or complex regulation [62, 81, 82]. All four datasets profile aerobic heterotrophic isolates on defined media; applicability to anaerobes and metagenome-assembled genomes is untested. All analyses rely on genomic annotations, which carry systematic biases across taxa [17, 83, 84]. Apparent concordance may therefore be lower in poorly studied lineages because of annotation gaps [85] rather than different biology, biasing the approach towards well-studied genera. Closing those gaps will require data from outside the annotation space, such as transcriptomic or regulatory profiles [86, 87], together with targeted characterisation of organisms outside the applicability domain, for which the low-confidence ranking of Figure 7 provides a starting point. Sequence-based language models offer a complementary route, since genomic and protein language models learn functional and regulatory relationships directly from sequence rather than from a curated ortholog vocabulary [88, 89], and protein-embedding classifiers have predicted bacterial phenotypes on taxa where database matching fails [90].

In summary, cross-dataset generalisation improved when training labels agreed with mechanistic predictions, but the resulting models are specialists that complement a curated pathway rule rather than replacing it, adding information mainly where the rule errs. Label–mechanism agreement is therefore a diagnosable and partially mitigable driver of shortcut learning, alongside sample volume and feature-space choice. Reliable genome-based phenotype prediction will therefore require curating training data on that criterion and reporting the domain in which a model applies. Which phenotypes are tractable at all, however, cannot be read from a model’s accuracy or its confidence, and must be established empirically. Extending the approach beyond well-characterised pathways will require a label-quality criterion that does not depend on a curated mechanistic predictor; phylogenetic agreement alone did not provide one, making ensemble disagreement and confidence-based label filtering [73] the natural candidates to test.

## Author Contributions

**D.K.**: Conceptualization, Data curation, Formal analysis, Investigation, Methodology, Software, Validation, Visualization, Writing – original draft, Writing – review & editing.

**P.R.**: Conceptualization, Formal analysis, Funding acquisition, Methodology, Software, Supervision, Writing – review & editing.

**C.N.**: Data curation, Software, Writing – review & editing.

**M.C.**: Data curation, Writing – review & editing. **Z.S.**: Data curation, Writing – review & editing. **W.J.R.**: Software, Writing – review & editing.

**M.P.J.**: Methodology, Writing – review & editing. **J.N.E.**: Data curation, Writing – review & editing. **J.P.F.**: Data curation, Writing – review & editing. **P.W.**: Data curation, Writing – review & editing. **M.B.C.**: Methodology, Writing – review & editing.

**D.A.P.**: Resources, Supervision, Writing – review & editing.

**R.W.C.**: Funding acquisition, Supervision, Writing – review & editing.

**C.S.H.**: Conceptualization, Funding acquisition, Supervision, Writing – review & editing.

**M.J.D.**: Funding acquisition, Supervision, Writing – review & editing.

**A.P.A.**: Conceptualization, Funding acquisition, Supervision, Writing – review & editing. **P.S.D.**: Conceptualization, Funding acquisition, Project administration, Supervision, Writing – review & editing.

P.R., C.S.H., M.J.D., A.P.A., and P.S.D. guided the analysis and substantially revised successive drafts of the manuscript.

## Acknowledgements

The authors thank the KBase team and members of the Plant-Microbe Interfaces Science Focus Area for discussions and infrastructure support.

## Use of artificial intelligence tools

Anthropic Claude (large language model) was used during manuscript preparation to assist with English-language editing of author-drafted text, to refactor analysis and figure-generation code from Jupyter notebooks into scripts, and to improve the visual aesthetics of matplotlib figures. OpenAI Codex (large language model) was used to audit journal-format compliance and assist with limited editing of author-drafted text. All code, data, scientific content, data interpretation, and final wording were reviewed and validated by the authors, who are responsible for the accuracy and integrity of the work.

## Supplementary Information

Supplementary Information (PDF) accompanies this paper: Supplementary Texts S1–S12 (dataset descriptions, genome quality, model selection, feature recovery, reliability, and data-volume analyses), Supplementary Figures S1–S7, and Supplementary Tables S1–S3.

## Data Availability

The four source datasets analysed in this study are cited under their own persistent identifiers: ATLeaf [36], Biolog [37], Marine [38], and Populus [39]; these records differ in whether they carry the phenotype measurements, the genomes, or both. The harmonised phenotype tables and genome accessions for all four datasets are therefore provided, together with the processed data, trained models, and figure source data generated during this study, in the Zenodo deposit [40]. Genome accessions and their source repositories (NCBI, JGI IMG, and BV-BRC) are listed in the deposited accession table, and the predicted proteomes of all genomes used for annotation are included in the Zenodo deposit. Computational genome annotations (RAST, KofamScan, GapMind) were produced using the cited tools on these genomes; dataset and genome-quality details are in Supplementary Texts S1–S2.

## Code Availability

The machine learning pipeline and Python library are available at https://github.com/ kbasecollaborations/trait-prediction. The manuscript source, analysis scripts, and figure generation code are available at https://github.com/kbasecollaborations/trait-prediction-manuscript.

## Supplementary Material

### Supplementary Text S1: Dataset Descriptions

We harmonised four experimental carbon source utilisation datasets spanning phylogenetically and ecologically diverse bacteria; analyses used the 15 phenotypes shared across all four datasets (Table S1). ATLeaf [1] tested *Arabidopsis thaliana* leaf-surface isolates on minimal agar medium. Biolog [2] was compiled from multiple published studies and unpublished contributions; growth calls derive from Biolog 96-well plate assays (GEN III, GN2 or Phenotype MicroArray, depending on the source), with a subset from other published growth-assay formats, and source studies for the published strains are given in the deposited metadata [3]. Marine [4] assessed heterotrophic marine bacteria. Populus [5] is a Biolog carbon-source assay of ORNL Plant-Microbe Interfaces isolates from the *Populus deltoides* rhizosphere and endosphere, recording optical density in MOPS minimal medium. Because of its small sample size and narrow phylogenetic breadth, Populus was excluded from selected feature-stability comparisons.

Generic carbon-source names such as “glucose” or “serine” do not specify stereochemistry, and the four datasets did not always assay the same isomer under the same name, so each phenotype was bound to one explicit source substrate, retaining the physiologically utilised isomer where a dataset assayed more than one. Identities were verified against Table S1 of Schäfer et al. [1] for ATLeaf, Supplementary Table 2 of Gralka et al. [4] for Marine, and the Biolog PM1 and PM2A plate maps for the Biolog and Populus assays. substrate identity common15.csv in the Zenodo deposit [3] records the resulting bindings, with a companion table covering all 240 carbon sources. Two cases warrant comment. The Marine *alanine* well contained DL-alanine, whereas the other three datasets assayed L-alanine; a racemate well scores positive if either enantiomer is utilised, so the Marine column is a union trait rather than a different trait. For *glucose*, the Populus alpha-D-glucose well was retained over the L-glucose well, which these taxa do not catabolise; the anomeric distinction is immaterial in solution because the free sugar interconverts by mutarotation, whereas for methyl glucosides and galactosides the locked acetal makes the two anomers separate traits taken up by different systems.

### Supplementary Text S2: Genome Quality Assessment

Genome quality was assessed with CheckM2 v1.1.0 [6] and recorded as a covariate rather than applied as an exclusion criterion, so that performance reflects the assemblies a user would realistically supply. Of the 755 modelled strains with a CheckM2 record, 689 met the MIMAG high-quality thresholds (completeness ≥90%, contamination ≤5%) and 732 met the medium-quality thresholds (completeness ≥50%, contamination ≤10%). Repeating the CatBoost KOFAM analysis for all 15 shared phenotypes with training, validation, and test sets each restricted to a quality tier left balanced accuracy essentially unchanged. Relative to the unfiltered data, the high-quality tier shifted it by a median of +0.004 under random splits (75 phenotype-fold pairs, Wilcoxon *p* = 0.03) and +0.015 under cross-dataset splits (58 pairs, *p* = 0.003), and the medium-quality tier by less than 0.001 (*p* = 0.72) and +0.005 (*p* = 0.05). These shifts are more than an order of magnitude smaller than the gap between random and cross-dataset splits, so genome quality filtering does not alter any conclusion reported here.

As an independent secondary audit, we measured pangenome completeness as the fraction of species-specific core genes detected by MMseqs2 [7] at 90% identity, 80% coverage, and E-value 10^−3^. 353 of the 394 assessed genomes with both a species assignment and a core-gene set (89.6%) exceeded 90% completeness (Figure S4).

### Supplementary Text S3: Model Selection and Evaluation Details

#### GapMind pathway annotation

Carbon-utilisation pathway completeness was computed with GapMind [8, 9] from the PaperBLAST suite (https://github.com/morgannprice/PaperBLAST, commit 1033662, accessed May 2025). For each pathway step, candidate genes were found by USEARCH v11.0.667 [10] against curated reference databases (minimum 30% identity) and by HMMER v3.3.1 profile search [11]. Reverse searches validated candidates, and required-step matches, alternative routes, and isofunctional enzymes determined the 0–2 pathway-completeness scores and the Methods thresholds.

#### Model family and hyperparameters

We compared gradient-boosted trees, regularised-linear and projection methods, Random Forest, and a phylogeny-aware model across 15 phenotypes and four evaluation regimes. 88% of phenotype–regime cells tied within 0.05 balanced accuracy, with boosted trees strongest and Random Forest the only consistent under-performer (Table S2).

This comparison used one protocol without early stopping, so its absolute values differ by ∼0.02 in balanced accuracy from the early-stopping CatBoost results. CatBoost was retained at 1000 iterations, learning rate 0.03, depth 4, L2 regularisation 15, 10% random feature subsampling, Bayesian bootstrap, and logloss early stopping with patience 50, because further phenotype-specific tuning did not improve accuracy. Figure 3C used a fixed 500-iteration CatBoost model without early stopping. The minority-class filter (Methods) was computed on the test subset plotted in each analysis and applied identically to both series in every machine-learning-versus-GapMind comparison (scripts/minority filter.py).

#### Replicate genomes and random-holdout overlap

Among the 755 KOFAM feature profiles entering at least one shared-phenotype split, 17 formed eight exact-duplicate groups, each confined to a single source dataset, so leave-one-dataset-out evaluation contained no exact-vector train–test overlap. Across random-holdout splits, approximately 1.4% of test predictions had an identical same-label vector in the training pool and approximately 9% a nearest training vector at Jaccard ≥ 0.98 (scripts/leakage check feature dup.py). Because both pre-processing filters are label-free, applying them across the full dataset rather than per split cannot leak labels, although the retained feature set reflects the pooled feature distribution; classifiers were still fitted on training samples only.

### Supplementary Text S4: Histidine Feature Comparison

Table S3 gives the histidine worked example under full-data and concordant training. Stable features ranked among the top 10 by mean absolute SHAP value in at least 70% of 20 runs, with cluster labels denoting SHAP-supervised redundancy groups (Methods). Under full-data training, the cluster containing K01468 and K01712 was already shared in all three comparisons (both KOs for ATLeaf and Biolog, K01712 alone for Marine), whereas K01745 and K05603 appeared only among dataset-specific stable features. Under concordant training, both K01468 (imidazolonepropionase) and K01712 (urocanate hydratase) were shared in all three comparisons, K05836 (histidine-utilisation repressor) became a shared feature in all three, the shared-cluster count rose from 4 to 8, the held-out-alone count fell from 6 to 3, and pathway concentration on map00340 rose from 17% to 33%. The example shared feature column of Table 1 shows a shared KO where the two models select the same feature outright, and otherwise a representative KO from each shared redundancy cluster; at most two are listed, so the column is illustrative rather than exhaustive, and it is empty only where no cluster is shared. Features are ranked by the number of held-out comparisons in which they recur, and those whose redundancy cluster touches the row’s KEGG map are listed first; off-pathway transporters or regulators appear only where no shared cluster touches the map (for glycerol, the *glpF* facilitator and the *glpR* repressor; for arginine, off-pathway choline/betaine and sarcosine-oxidase clusters). The displayed features do not enter the (% in pathway) statistic, which scores canonical pathway-map residency across all shared clusters rather than the one or two shown.

### Supplementary Text S5: Concordant Training Under the KOFAM Feature Space

GapMind concordance selected training samples, whereas KOFAM annotations supplied model features, decoupling the selection rule from the feature space. Under the same KOFAM feature space, concordant training improved cross-dataset balanced accuracy on concordant test samples for most phenotype–split combinations (Figure S5). The feature-recovery analysis nevertheless remains biologically related to the selection rule, since GapMind and KOFAM annotate the same genomes. However, GapMind scores curated reference proteins rather than KEGG modules, SHAP stability across datasets is not entailed by the selection rule, and several phenotypes recovered predominantly off-map features. A size- and class-matched random-subset control recovered 0.38 ± 0.03 shared clusters (mean ± SD across ten draws), similar to full-data training (0.5) and below concordant training (1.3), indicating that the stability gain was specific to concordance rather than sample count (scripts/figure5/figure5b random control.py).

Raw GapMind features without correlation filtering were used for the rule-reconstruction reference (Figure 5A) and the phenotype-filtered comparison (Figure 6D) because the standard 0.95 correlation filter collapses repeated phenotype-prefixed representations of the same gene and can deplete individual pathway feature sets. Because these features are the evidence the concordance rule itself is computed from, this reference bounds how far that rule can be reconstructed by the ML model from its own inputs, and is not an estimate of annotation quality.

Feature selection within the combined matrix indicates why concatenating annotation types did not improve transfer (Figure 1C). RAST subsystem terms contribute 62.6% of its 17,152 correlation-filtered features, KOFAM 31.8%, and GapMind only 5.7%; yet among the top ten features by mean absolute SHAP value across the cross-dataset splits, GapMind terms account for 43.0% of selections, a 7.6-fold enrichment, RAST terms for 26.8%, a 0.4-fold depletion, and KOFAM terms in proportion to their share. The comprehensive set therefore inherits the same dependence on curation as the phenotype-filtered set without its accuracy: the roughly 34 curated features of a single phenotype outperformed all 17,152 combined features under cross-dataset evaluation (Figure 6D).

### Supplementary Text S6: Learning Curves for the Shared Phenotypes

For most of the 15 shared phenotypes, concordance-trained KOFAM performance approached its all-available-sample value within 50 to 100 training samples (Figure S6). Under cross-dataset evaluation, 11 of 15 phenotypes reached 90% of their all-sample balanced accuracy with 50 samples and 14 of 15 with 100, fructose alone requiring 200. Median performance at 50 samples was 93.6% of the all-sample value for concordant training and 91.0% for full-data training. Early saturation does not by itself indicate a well-predicted phenotype. The five phenotypes saturating with 25 cross-dataset samples had a median all-sample balanced accuracy of 0.63, against 0.73 for the remainder, and cross-dataset gains between 25 samples and the full set were larger for phenotypes that ultimately performed better (Spearman *ρ* = 0.78, *p* = 0.0007). Glucose, for example, saturated immediately at a balanced accuracy of 0.51. Additional samples therefore helped most where transfer was already succeeding, consistent with a representational rather than a statistical limit on the phenotypes that failed to transfer.

### Supplementary Text S7: SHAP-Supervised Redundancy Clustering Procedure

Stable features were identified with a fixed 500-iteration CatBoost model trained without early stopping, using 20 random seeds per phenotype–dataset configuration. A feature was designated stable when it ranked among the top ten by mean absolute SHAP value in at least 70% of those runs. For the concordance-trained comparison in Figure 5B, each configuration was first screened to 300 candidate KOFAM features using CatBoost PredictionValuesChange importance, after which the 20-seed SHAP stability analysis was applied to the candidate set.

Because correlated KOs can represent the same predictive signal, the stable-feature comparisons in Figures 4C and 5B intersect clusters rather than raw KOs. For each phenotype, the union of stable KOs was clustered on the pooled concordant feature matrix across all four datasets with shap.utils.hclust in SHAP v0.50.0 [14–16]. Supplied with the phenotype labels, the routine fitted a univariate XGBoost model of the phenotype from each KO, then a second model predicting that first-stage output from every other KO; each directional distance was 1 − *R*^2^. The smaller directional distance for each KO pair formed a single-linkage matrix, which was cut at a distance threshold of 0.5.

For alanine, mannose, and serine, stable features from different concordance-trained models did not enter the same cluster but remained correlated within some held-out datasets (for mannose, transporters K10441 and K10545 at |*r* | = 0.67 in concordant ATLeaf samples). These features were heterogeneous carbohydrate-utilisation and accessory genes rather than one shared pathway, indicating a broader, less substrate-specific mode of transfer; for alanine, GapMind’s pathway definition encodes only transport, so concordance reflects uptake rather than catabolic coverage.

### Supplementary Text S8: Reliability and Experiment-Prioritisation Details

CatBoost probabilities were not post hoc calibrated, so confidence ranked predictions only. Expected calibration error used ten equal-width bins, pooled across phenotypes for each model. Neither mean confidence nor the fraction of predictions (with confidence of at least 0.8) predicted per-phenotype cross-dataset balanced accuracy across the 15 phenotypes (Spearman *ρ* between −0.18 and −0.05, both *p* > 0.5).

Feature-space novelty was the mean Jaccard distance from a held-out genome to its five nearest training genomes in KOFAM space. The high-novelty strategy took the candidates ranked highest on this measure; the diversity strategy took candidates evenly spaced along the same ranking. Each eligible held-out dataset was split approximately 50:50 into candidate and evaluation pools, stratified where possible, and 25 candidate labels were added to the concordant training set. Models used 120 CatBoost iterations, depth 4, and learning rate 0.05 without evaluation-set early stopping, across three seeds and four held-out datasets. Because the random baseline also adds labels from the held-out dataset, it carries the benefit of initial exposure, and the other strategies are credited only with improvement beyond it. After averaging seeds within 23 phenotype–held-out-dataset cells, low-confidence selection exceeded random selection (*p* = 0.024, Benjamini–Hochberg-adjusted *q* = 0.052); across the six phenotype means, four favoured low-confidence selection and the paired test was not significant (*p* = 0.156).

### Supplementary Text S9: Composite Confidence Weighting

The composite score used to weight training and validation samples (Figure 6B) was calculated as follows:

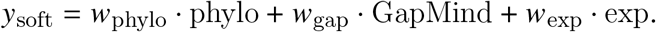

Within each split, the phylogenetic component was the mean experimental label among the *k* = 3 nearest training genomes and therefore took values 0, 1/3, 2/3, or 1. The training genome was excluded from its own neighbours. The experimental component was the binary phenotype label used only for training and validation-set quality assessment. GapMind categories mapped to six numeric levels: complete, 1.0; likely complete, 0.9; steps missing medium, 0.6; steps missing low, 0.3; not present, 0.1; and no evidence or uncategorized, 0.0. After clipping *y*_soft_ to [0.01, 0.99], every scored sample was retained and assigned the recall-prioritised weight

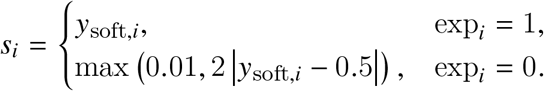

The asymmetric weighting prioritised recall while ensuring that every phylogenetic conflict reduced confidence. In the mechanism-free setting, complete phylogenetic conflicts remained in training with weights 0.60 for positive labels and 0.20 for negative labels, while complete agreements received 0.99 and 0.98, respectively. Four weight configurations (*w*_phylo_, *w*_gap_, *w*_exp_) were evaluated: the mechanism-free (0.40, 0.00, 0.60) and three GapMind-weighted settings (0.20, 0.30, 0.50), (0.15, 0.40, 0.45), and (0.10, 0.50, 0.40). All weights were applied to training and validation samples only; held-out test labels were not used to calculate them, and held-out test samples were unchanged. Only genomes placed in the phylogenetic tree could be scored, so the mechanism-free analysis retained 80% of samples as rows (*n* ≈ 427; summed sample weight ≈ 374), compared with *n* ≈ 534 without filtering and *n* ≈ 385 under concordance filtering.

#### Performance across the weight sweep

Across the four settings, mean cross-dataset balanced accuracy on the full held-out test set varied by less than 0.005, while mean recall rose from 0.759 at *w*_gap_ = 0 to 0.826 at *w*_gap_ = 0.5 as precision declined from 0.733 to 0.701. At *w*_gap_ = 0.5 recall exceeded GapMind by a mean of 0.089 (*p* = 0.010) but precision was lower by 0.066 (*p* = 0.010) and specificity by 0.198 (*p* < 0.001), so the endpoint shifted the operating point towards positive calls rather than adding discriminative information. Mean balanced accuracy was 0.68 for concordant training and 0.644 for mechanism-free weighting (paired Wilcoxon *p* = 0.23), against 0.646 without filtering and 0.69 for GapMind. Mechanism-free weighting increased mean recall by 0.011 over unfiltered ML (higher for 12 of 15 phenotypes, *p* = 0.11) while reducing mean specificity by 0.015. Against GapMind on the same held-out genomes, concordant training recovered significantly more true positives for 9 of 15 phenotypes and fewer for 4, whereas mechanism-free weighting was higher for 6 and lower for 6; neither mean recall shift was significant (*p* = 0.095 and *p* = 0.56, respectively).

### Supplementary Text S10: Per-Phenotype Detail for Concordant-Training Analysis

#### GapMind-feature glucose gap

The largest feature-resolution gap between GapMind and KOFAM feature models was glucose, for which GapMind-feature and KOFAM cross-dataset balanced accuracy were 0.77 and 0.51, respectively. GapMind uses substrate-specific transporter steps such as MFS-glucose, SSS-glucose, and ptsG-crr that lack direct KOFAM equivalents.

#### Discrimination within a GapMind call

Rescue rates, the fractions of GapMind errors on the discordant cross-dataset subset that the concordance-trained model classified correctly, appeared higher for false negatives than false positives (42% versus 20%, paired Wilcoxon *p* = 0.018), but the difference vanished once each phenotype’s marginal model-call rate was subtracted (−0.19 versus −0.19, *p* = 0.98), because a hard-call rescue rate is bounded by how often the model emits that class at all. We therefore scored discrimination within each GapMind call by the predicted growth probability, which is threshold-free and on which GapMind scores 0.5 by construction, its call being constant within a stratum. The concordance-trained model separated growers from non-growers in both strata (pooled AUC 0.674 among GapMind-negative and 0.676 among GapMind-positive genomes; per-phenotype means 0.636 and 0.656), but improved on full-data training only in the negative stratum (0.636 versus 0.534, higher for 13 of 15 phenotypes, *p* < 0.001, Benjamini– Hochberg *q* = 0.003; positive stratum 0.656 versus 0.646, 8 of 15, *p* = 0.76). This matches the expected genomic basis of the two error types: growth despite an incomplete annotated pathway leaves genomic content that concordant training sharpens, whereas a complete but unexpressed pathway does not. Three phenotypes fell below 0.5 in the GapMind-negative stratum (glycerol 0.455, arginine 0.443, serine 0.258).

#### What the recovered genomes carry

False-negative growers used more carbon sources than GapMind-negative non-growers (9.5 versus 5.2 of the remaining 14; Mann–Whitney *p* < 10^−200^) and encoded more sugar-ABC transport components (6.5 versus 4.3; *p* < 10^−60^). The features concordant models used to predict these genomes were dominated by broad carbohydrate transporters and regulators, and in 9 of 15 phenotypes a transporter rather than an enzyme was the GapMind step most often absent, supporting a metabolic-generalist rather than substrate-specific interpretation. Candidate exceptions with greater substrate-specific plausibility included sucrose uptake through a broad multiple-sugar ABC importer (K02025) and the arabinonate dehydratase K26399, a member of the D-altronate-dehydratase family [17, 18], but neither establishes a validated alternative route. Under balanced class weights, retaining these false negatives in training rather than filtering them out increased recall by 0.030 but reduced specificity by 0.088 and balanced accuracy by 0.029 relative to concordance filtering, so it is preferable only when recall is prioritised (scripts/figure5/fp only filter.py). Partitioning false negatives by the genomic presence of their catabolic machinery separated a recoverable fraction from a larger residual lacking such support, which was enriched for confident model–GapMind agreement against the experimental label (odds ratio 3.3), which could be due to experimental mislabelling (Supplementary Text S12).

#### Enforcing concordance at prediction time

Concordance with GapMind can be enforced at two different points in the pipeline. Concordant training, used throughout the main text, filters the training set: it retains the roughly 70% of training samples whose labels agree with GapMind and then predicts for every new genome from KOFAM features alone, with no GapMind call needed at prediction time. Alternatively, concordance can be enforced on the predictions instead: a model trained on all samples predicts for every genome, but only the predictions that agree with that genome’s GapMind call are retained, and the rest are withheld. On the pooled held-out test sets this retained 72% of predictions and reached cross-dataset balanced accuracy 0.77 on the retained subset. Because the model abstains on the remaining 28% of genomes, this value describes only the retained predictions and is not directly comparable with performance measured over a full evaluation set. Prediction-time agreement therefore buys higher accuracy on the predictions it retains at the cost of making no prediction for the rest, and it requires a GapMind prediction for every genome at deployment, whereas concordant training requires GapMind only once, when the training set is assembled.

### Supplementary Text S11: Adding Data Volume Versus Concordance Filtering

Because the manuscript dataset contained up to 780 feature-linked strains, we tested whether added training volume could substitute for concordance-based curation [19, 20]. CatBoost classifiers using KOFAM features were evaluated on the full labels of each held-out manuscript dataset across 13 phenotypes, four leave-one-dataset-out splits, and five seeds, training on concordant samples from the other three datasets (C), all samples from those datasets (F), or F plus BacDive (FB). The BacDive input was a fixed snapshot of 8,501 genomes with KOFAM annotations [21]. After label–feature intersection and removal of accessions overlapping the held-out test set, up to approximately 1,800 BacDive genomes per phenotype entered training, with balanced class weights applied uniformly because BacDive labels were strongly positive-skewed.

FB improved balanced accuracy over F (Figure S7A; 0.725 versus 0.677, paired Wilcoxon *p* < 0.001, higher for 12 of 13 phenotypes) but was not significantly different from C (0.725 versus 0.706, *p* = 0.20; Matthews correlation coefficient *p* = 0.53; area under the ROC curve *p* = 0.99), with higher precision offset by lower recall. BacDive alone matched the concordant set (0.709 versus 0.706, *p* = 0.53), whereas adding BacDive to the concordant set exceeded it (0.729 versus 0.706, *p* = 0.03 across the 45 phenotype–held-out cells, *p* = 0.09 across the 13 phenotype means). Coherence and volume are therefore complementary rather than alternatives: concordance filtering reached comparable cross-dataset performance from roughly four-fold fewer training samples, and added volume improved on it further only when applied to the already-filtered set.

Feature selection was less reproducible as volume increased (Figure S7B,C): mean pairwise top-10 Jaccard similarity was 0.378 for C, 0.316 for F, and 0.253 for FB (C versus FB, *p* < 10^−8^), and concordant models more consistently selected committed-pathway features, although mannose was an exception. C used fewer samples and produced more reproducible, mechanism-aligned feature rankings, a pattern consistent with reduced shortcut learning [22].

### Supplementary Text S12: Cross-Dataset Label Inconsistency Among Near-Identical Strains

At phylogenetic distance *d* ≤ 0.01, phenotype-specific comparisons between near-identical strains conflicted in 28.0% of cross-dataset cases (73 of 261) and 10.1% of within-dataset cases (782 of 7,706; odds ratio 3.44), with a monotone excess at stricter thresholds. Pair-level logistic adjustment for dataset positive-call rates reduced but did not remove the association (adjusted odds ratio 2.69; 1.86 conditioning on both datasets’ rates), and because comparisons reuse strains across pairs and phenotypes, these odds ratios are descriptive, with independence-based confidence intervals and *p*-values not interpreted. The analysis cannot distinguish assay differences, ecological or strain-history divergence, accessory-gene or regulatory differences, and curation effects, and does not show that concordance filtering specifically identifies the conflicting pairs.

---

**Figure S1:**
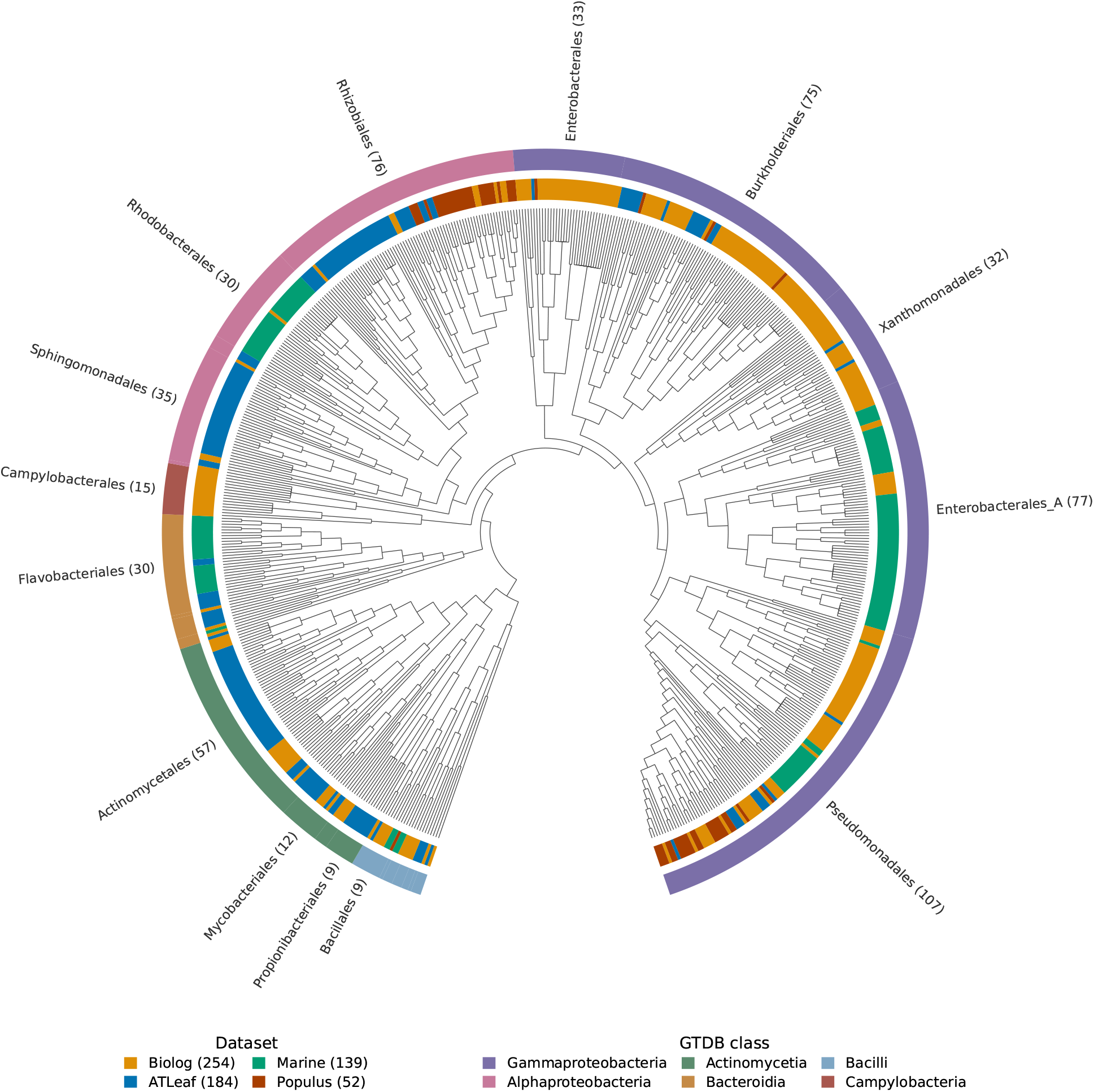
The four datasets span overlapping but unevenly sampled regions of the bacterial tree. Of the 819 input strain records, 629 GTDB-placed tips were retained in the pruned Genome Taxonomy Database reference tree [12], with distances calculated using ETE 3 [13]. The tree is drawn as a cladogram, so branch lengths are not to scale. Phylogenetic distances are reported in Figure S2. The inner ring denotes dataset membership and the outer bar the GTDB order, coloured by GTDB class. Orders with at least eight tips are labelled with their tip count, and tip labels are omitted for legibility.

**Figure S2:**
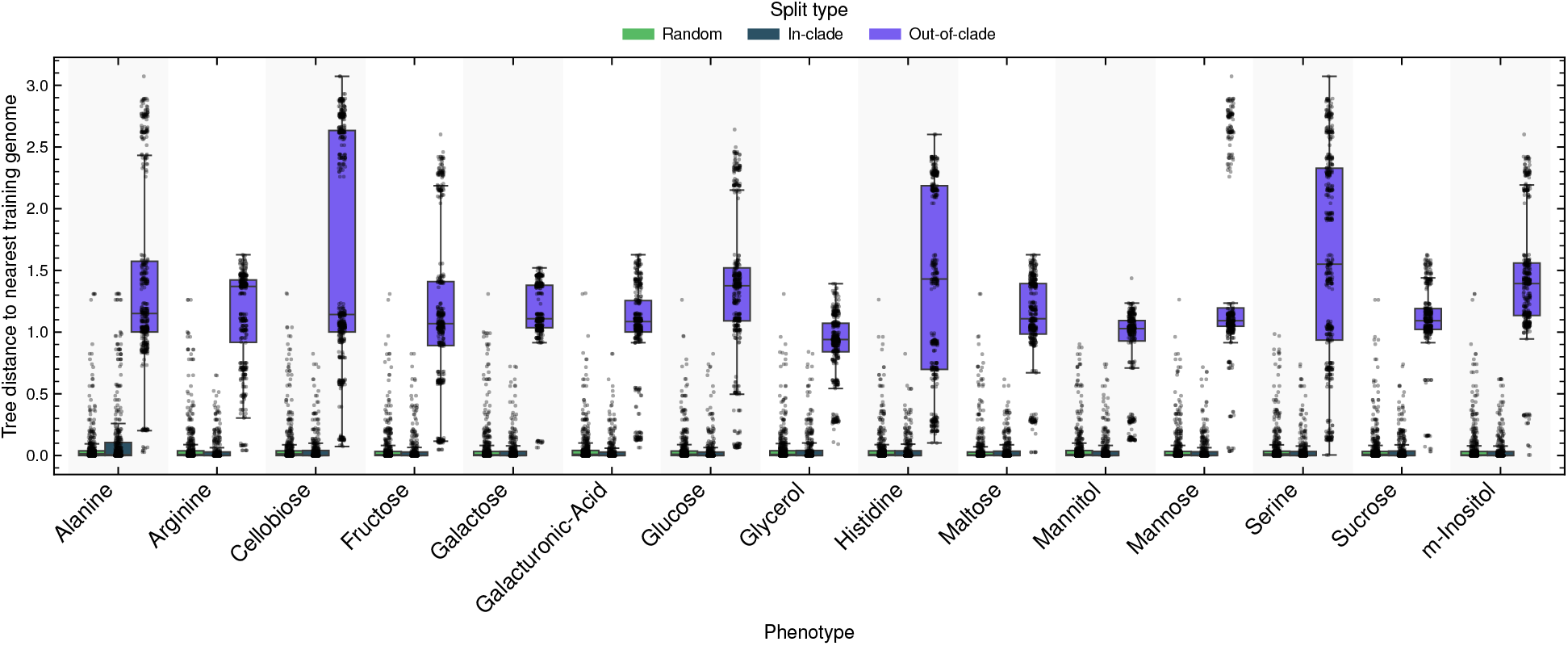
Only out-of-clade splits place test genomes far from their training set. Tree distance (summed branch lengths) from each held-out test genome to its nearest training genome under random-holdout, in-clade, and out-of-clade splits, pooled across five splits per phenotype for the 15 shared phenotypes. Boxes are coloured by split type (random, green; in-clade, dark teal; out-of-clade, purple) and show the median and interquartile range; points are individual test genomes (random *n* = 8,289; in-clade *n* = 5,595; out-of-clade *n* = 6,212). Random-holdout and in-clade test genomes lie close to the training set (medians 0.009 and 0.000), whereas out-of-clade splits produce much larger distances (median 1.109), validating the intended separation of the evaluation regimes in Figure 3.

**Figure S3:**
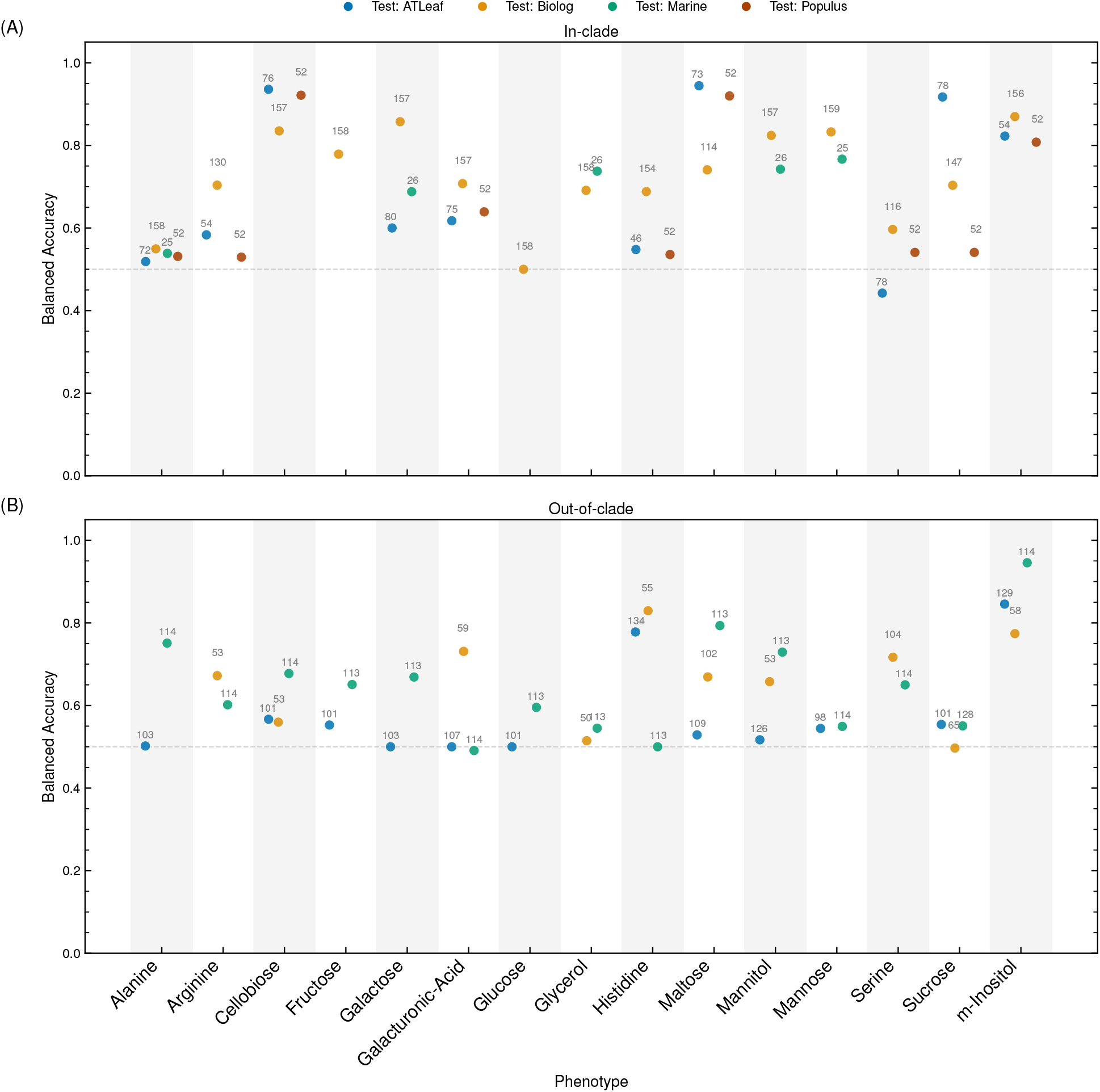
Cross-dataset accuracy is lower for genomes without close training relatives. CatBoost balanced accuracy using KOFAM features across four leave-one-dataset-out combinations. (**A**) In-clade, test samples sharing a cluster with at least two training samples; (**B**) out-of-clade, test samples sharing a cluster with fewer than two, using the Figure 3C clustering procedure. Points are coloured by held-out test dataset. Cells with fewer than 10 test or minority-class samples were excluded (44 of 120, including all Populus-held-out out-of-clade cells). Mean out-of-clade balanced accuracy was lower than in-clade accuracy (0.621 versus 0.701). Numerals give test-subset sizes.

**Figure S4:**
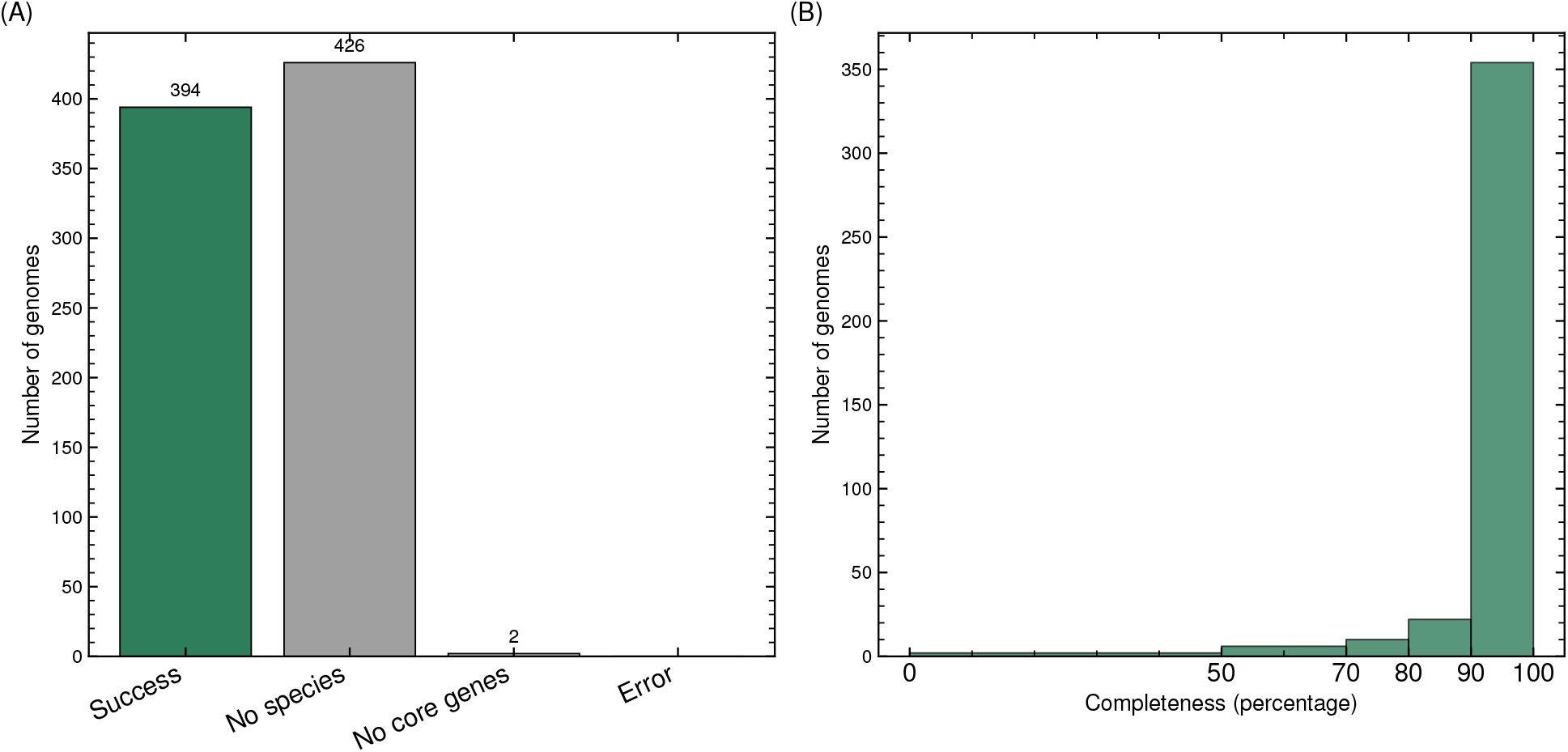
Pangenome completeness was assessed for a subset of genomes, nearly all of which were near-complete. Completeness was the fraction of species core genes detected by MMseqs2 at 90% identity, 80% coverage, and E-value 10^−3^. (**A**) Assessment status. (**B**) Completeness among successfully assessed genomes, most of which exceeded 90%.

**Figure S5:**
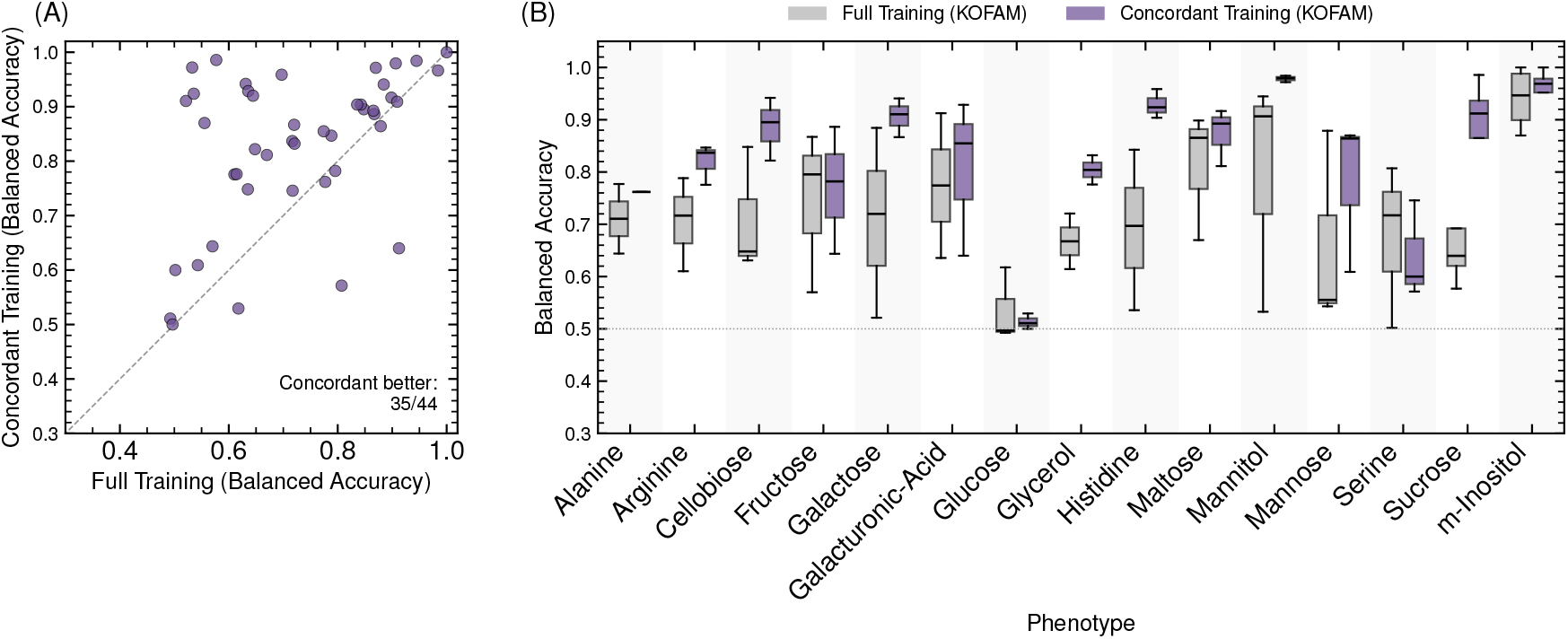
Concordant training improves cross-dataset generalisation under KOFAM features. (**A**) Cross-dataset balanced accuracy of concordance-trained versus full-data KOFAM models on concordant test samples; each point is one phenotype–split combination (*n* = 44), and the annotation gives the number of combinations on which concordant training was better. (**B**) Per-phenotype distributions for full-data (grey) and concordant (purple) training; boxes show the median and interquartile range, and the dotted line marks the 0.5 chance level. Each box summarises the one to four cross-dataset splits available for that phenotype, so boxes for phenotypes with a single split appear as a line.

**Figure S6:**
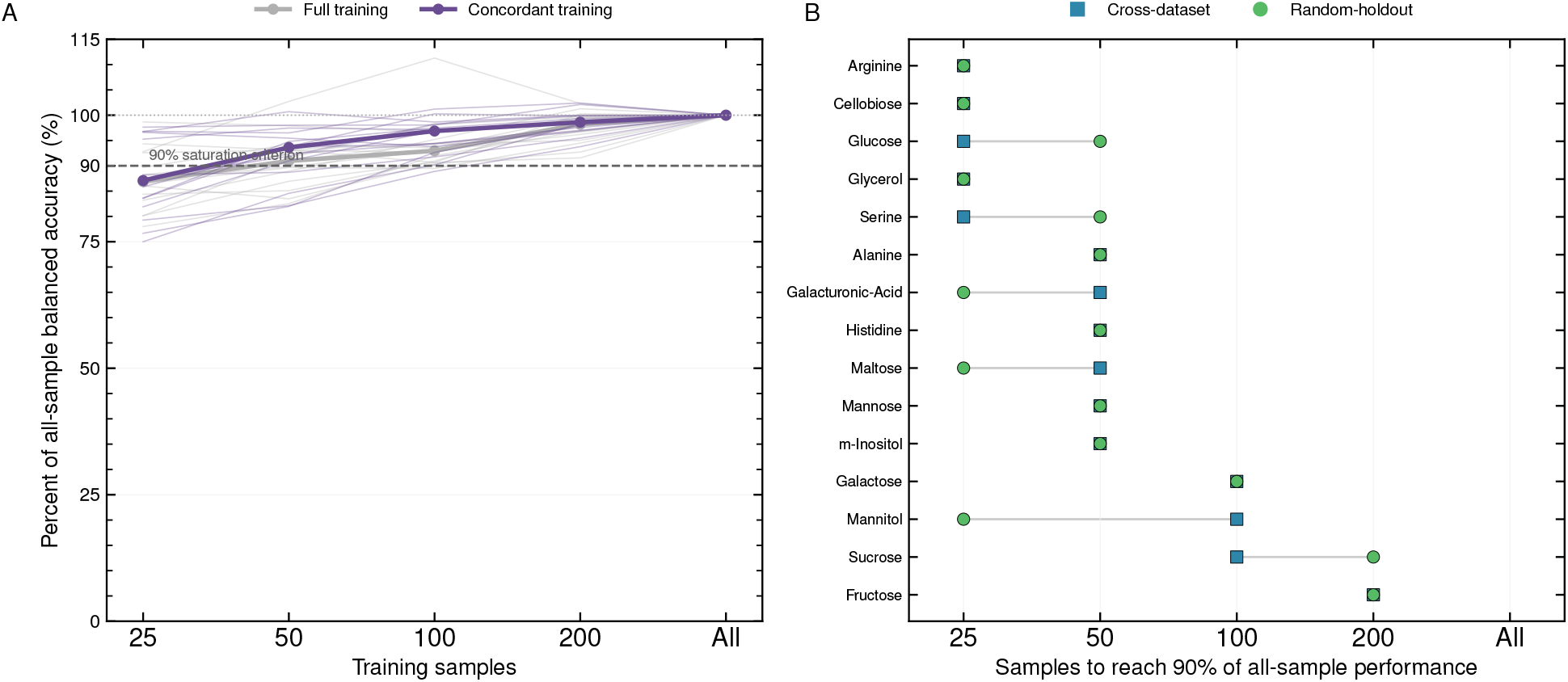
Cross-dataset performance saturates at 50–100 training samples for most phenotypes. (**A**) Cross-dataset balanced accuracy at each training-set size, expressed as a percentage of the same phenotype’s all-sample value, for full-data (grey) and concordant (purple) training. Thin lines are individual phenotypes and bold lines their medians; the dashed line marks the 90% saturation criterion and the dotted line the all-sample value. Values above 100% arise where an intermediate training-set size outperformed the full set. (**B**) Smallest training-set size at which each phenotype reached 90% of its all-sample performance under cross-dataset (blue squares) and random-holdout (green circles) evaluation, ordered by the cross-dataset value. Points are means across repeats and across the splits available for each phenotype, at requested sizes of 25, 50, 100, 200, and all available samples.

**Figure S7:**
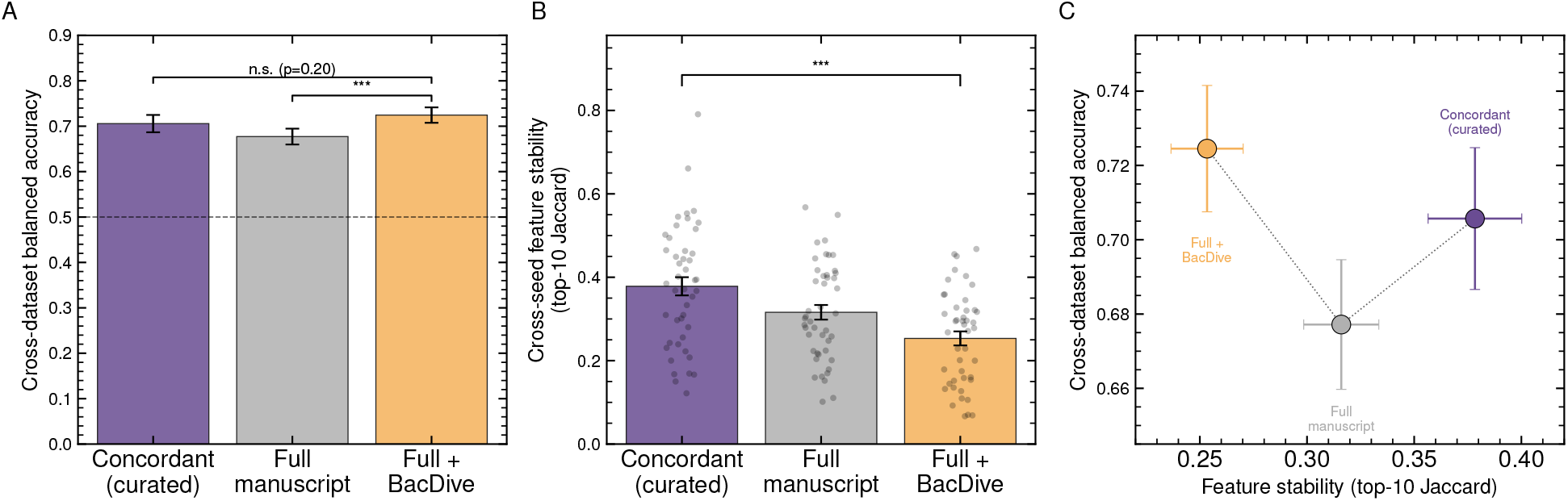
Added data volume substitutes for curation on accuracy, but not on feature reproducibility. CatBoost classifiers using KOFAM features were tested on each held-out manuscript dataset under C (concordant), F (full manuscript), and FB (full manuscript plus BacDive [21]) training. (**A**) Cross-dataset balanced accuracy per strategy; the dashed line marks chance (0.5). (**B**) Cross-seed feature stability, the mean pairwise Jaccard of the top-10 CatBoost-importance features across the five seeds; points are individual phenotype–held-out sets. (**C**) Per-strategy cross-dataset balanced accuracy versus feature stability. Bars and markers show means, error bars show the standard error across 45 phenotype–held-out cells, and brackets show paired Wilcoxon tests. “n.s.” denotes not significant and *** denotes *p* < 0.001.

**Table S1:** The four datasets differ in size, growth assay format, and positive-call rate. “Original” gives the source-dataset size; “Retained Phenotype Records” gives the records with an available genome assembly; “Feature-linked” gives phenotype records with matching processed genomic features. “Positive-Call Rate” is the fraction of non-missing labels scored as growth across the 15 shared phenotypes, the quantity adjusted for in the label-conflict analysis (Supplementary Text S12). All datasets entered aggregated and leave-one-dataset-out analyses; Populus was excluded only from selected feature-stability comparisons.

| Dataset | Original Strains | Original Carbon Sources | Retained Phenotype Records | Feature-linked Strains | Shared Phenotypes | Positive-Call Rate | Growth Assay |
| --- | --- | --- | --- | --- | --- | --- | --- |
| ATLeaf | 224 | 45 | 206 | 206 | 15 | 0.47 | Agar plate |
| Biolog | 362 | 64 | 362 | 362 | 15 | 0.65 | Biolog plates and related assays |
| Marine | 175 | 100 | 172 | 157 | 15 | 0.51 | Liquid culture OD |
| Populus | 58 | 190 | 55 | 55 | 15 | 0.74 | Liquid culture OD |

**Table S2:** Model family has little effect on accuracy. Mean balanced accuracy across the 15 shared phenotypes under random-holdout and leave-one-dataset-out evaluation on full and GapMind-concordant subsets. All models used KOFAM features and one protocol without early stopping; per-cell standard deviation was approximately 0.05–0.10, and additional families are in the deposited analysis.

| **Model** | **Full data** |  | **Concordant** |  |
| --- | --- | --- | --- | --- |
|  | **Random** | **Cross-dataset** | **Random** | **Cross-dataset** |
| CatBoost | 0.829 | 0.633 | 0.957 | 0.871 |
| LightGBM | 0.836 | 0.638 | 0.961 | 0.887 |
| Elastic-Net | 0.829 | 0.643 | 0.945 | 0.827 |
| Random Forest | 0.823 | 0.612 | 0.933 | 0.775 |

**Table S3:** Concordant training stabilises the same histidine-degradation cluster across datasets. Shared features occur in both the three-dataset and held-out-alone models; unique features occur only in the held-out-alone model. Cluster labels mark SHAP-supervised redundancy groups, and module coverage counts one representative per cluster across the three comparisons. Features appeared in at least 70% of 20 training runs. Table 1 and Figure 5B count clusters rather than KOs.

| Training set | Held-out dataset | Shared stable features | Unique to held-out-alone model |
| --- | --- | --- | --- |
| Full data | ATLeaf | [Cluster C] K01468: imidazolonepropionase [EC:3.5.2.7]<br>[Cluster C] K01712: urocanate hydratase [EC:4.2.1.49] | K00547: homocysteine S-methyltransferase [EC:2.1.1.10]<br>[Cluster B] K01032: 3-oxoadipate CoA-transferase, beta subunit [EC:2.8.3.6]<br>[Cluster C] K01745: histidine ammonia-lyase [EC:4.3.1.3]<br>K02623: LysR family transcriptional regulator, pca operon transcriptional activator<br>[Cluster A] K05603: formimidoylglutamate deiminase [EC:3.5.3.13]<br>K09975: uncharacterized protein |
|  | Biolog | [Cluster C] K01468: imidazolonepropionase [EC:3.5.2.7]<br>[Cluster C] K01712: urocanate hydratase [EC:4.2.1.49] | K00433: non-heme chloroperoxidase [EC:1.11.1.10] |
|  | Marine | [Cluster C] K01712: urocanate hydratase [EC:4.2.1.49]<br>K09975: uncharacterized protein | None |
| Module coverage (M00045 *Histidine degradation*): shared 1/2 (50%); unique 2/7 (29%) |  |  |  |
| Concordant | ATLeaf | [Cluster C] K01468: imidazolonepropionase [EC:3.5.2.7]<br>[Cluster C] K01712: urocanate hydratase [EC:4.2.1.49]<br>K05836: GntR family transcriptional regulator, histidine utilization repressor | [Cluster B] K00481: p-hydroxybenzoate 3-monoxygenase [EC:1.14.13.2]<br>[Cluster B] K01032: 3-oxoadipate CoA-transferase, beta subunit [EC:2.8.3.6]<br>[Cluster A] K01458: N-formylglutamate deformylase [EC:3.5.1.68]<br>[Cluster C] K01745: histidine ammonia-lyase [EC:4.3.1.3]<br>[Cluster A] K05603: formimidoylglutamate deiminase [EC:3.5.3.13]<br>K09975: uncharacterized protein |
|  | Biolog | [Cluster C] K01468: imidazolonepropionase [EC:3.5.2.7]<br>[Cluster C] K01712: urocanate hydratase [EC:4.2.1.49]<br>K05836: GntR family transcriptional regulator, histidine utilization repressor | [Cluster B] K02624: IclR family transcriptional regulator, pca regulon regulatory protein |
|  | Marine | [Cluster C] K01468: imidazolonepropionase [EC:3.5.2.7]<br>[Cluster C] K01712: urocanate hydratase [EC:4.2.1.49]<br>K05836: GntR family transcriptional regulator, histidine utilization repressor<br>K09975: uncharacterized protein | None |
| Module coverage (M00045 *Histidine degradation*): shared 1/3 (33%); unique 2/4 (50%) |  |  |  |

